# Evaluation of Connectivity Map shows limited reproducibility in drug repositioning

**DOI:** 10.1101/845693

**Authors:** Nathaniel Lim, Paul Pavlidis

## Abstract

The Connectivity Map (CMap) is a popular resource designed for data-driven drug repositioning using a large transcriptomic compendium. However, evaluations of its performance are limited. We used two iterations of CMap (CMap 1 and 2) to assess their comparability and reliability. We queried CMap 2 with CMap 1-derived signatures, expecting CMap 2 would highly prioritize the queried compounds; success rate was 17%. Analysis of previously published prioritizations yielded similar results. Low recall is caused by low differential expression (DE) reproducibility both between CMaps and within each CMap. DE strength was predictive of reproducibility, and is influenced by compound concentration and cell-line responsiveness. Reproducibility of CMap 2 sample expression levels was also lower than expected. We attempted to identify the “better” CMap by comparison with a third dataset, but they were mutually discordant. Our findings have implications for CMap usage and we suggest steps for investigators to limit false positives.

## Introduction

Connectivity Map (CMap) is a resource created to enable data-driven studies on drug mode-of-action and drug repositioning (Lamb et al., 2006). CMap works by comparing genes from user-provided “gene hit lists” (also called “signatures”) to a large reference gene differential expression (DE) database generated by perturbing cell lines with a wide range of chemical compounds (Subramanian et al., 2005). The outcome is a ranking of compounds that yield expression patterns most similar to that of the query hit list. Many researchers have applied CMap with goals ranging from identification of candidate drugs for repurposing to investigating the biological activity of drugs (Musa et al., 2018; Qu and Rajpal, 2012). In this paper we consider the question of whether CMap is an effective tool for this purpose.

Released in 2006 (Lamb et al., 2006), CMap version 1 (“CMap 1”) applied 1,309 compounds to five different cell lines at varying doses, yielding a corpus of 6,100 DE profiles. The Connectivity Map project recently entered a second iteration (“CMap 2”) as part of NIH’s Library of Integrated Network-Based Cellular Signatures (LINCS) program (Keenan et al., 2018; Subramanian et al., 2017). CMap 2 (also known as LINCS-L1000) contains 591,697 profiles generated from 29,668 compounds and genetic modifications (collectively called “perturbagens” in LINCS parlance) across 98 different cell lines. The massively increased scale and scope of CMap 2 raises prospects for improved pharmacogenomics studies. The CMap team also encourages researchers to use CMap 2 for ongoing research (https://clue.io/cmap).

CMap 2 differs in important technical ways from CMap 1. CMap 2 replaced the Affymetrix GeneChips used in CMap 1 with Luminex bead arrays that assay the expression of 978 “landmark” genes, with expression levels of another 11,350 genes inferred computationally (compared to 12,010 genes directly assayed in CMap 1). These differences raise the natural question as to whether CMap 2 is more reliable, or even comparable, to CMap 1. In the original publication introducing CMap 2, while the authors demonstrated agreement of the Luminex technology to RNA-sequencing, no comparison to CMap 1 was described (Subramanian et al., 2017). To our knowledge, there has been no detailed evaluation of the CMaps; there have been some forms of comparisons which further motivates the need for such an evaluation. Recently, Niepel et al. assessed the reproducibility of drugs effects on cell growth across LINCS centres, though they did not examine gene expression (Niepel et al., 2019). More directly relevant to our work, Iwata et al. reported that the gene expression profiles between the two CMap versions were poorly correlated despite having comparable data quality (Iwata et al., 2017). Their comparison however did not control for differences in compound concentrations, raising questions on the fairness of the comparison; neither did they test comparability of candidate drug recommendation results. In an anecdotal report, Zador et al. found that the candidate drug azacitidine was recommended by CMap 2 while being rejected by CMap 1 (Zador et al., 2018). On the other hand, there are also anecdotal reports of positive results, in which researchers have experimentally verified the biological effects of candidate drugs recommended by CMap 1 (Braconi et al., 2009; Brum et al., 2015, 2018; Byun et al., 2019; Johnstone et al., 2012; Luo et al., 2019; Schanstra et al., 2019; Wang et al., 2019; Wu et al., 2019) or CMap 2 (Ferguson et al., 2018; Leung et al., 2019; Manzotti et al., 2019; Ryals et al., 2018). An attempt to address the reproducibility question was presented by De Abrew et al., in which they generated an independent CMap-style dataset for 34 compounds in four cell lines (De Abrew et al., 2016). They compared compound retrieval performance to CMap 1 for the 17 compounds shared with their study (their work predates CMap 2). While they reported that the retrieval performance was generally positive, there was apparently a high degree of variability in performance, and no quantitative assessment was presented. They also suggested that agreement with CMap 1 was highest for the highest concentrations of compounds.

In this paper, we revisit the question of comparability and reliability of CMap 1 and 2. We demonstrate that candidate compound prioritization is generally very different between the two. We show that this is underpinned by the fact that the DE profiles for the same condition are generally poorly correlated both between and within CMap 1 and 2. We also show that reproducibility of DE profiles is predicated on DE strength, which is in turn influenced by compound concentration and cell line responsiveness. We also demonstrate that within-CMap 2 agreement of sample gene expression levels is poor, and acts as another predictor of DE reproducibility. We quantitatively re-evaluate the data of De Abrew et al. and show that their data also tend to disagree strongly with both CMap 1 and 2, even for the same compound concentration and cell line. Our results have implications for drug repurposing projects using either CMap version.

## Methods

Except where noted, analyses were performed using R and Python. Scripts and relevant data files are provided on Github (https://github.com/PavlidisLab/CMap-Evaluation).

### Processing of CMap 1 data

We used build 02 of CMap 1. The metadata was obtained from the Broad Institute’s CMap webpage (https://portals.broadinstitute.org/cmap/), while the differential expression data was obtained from the Broad Institute’s FTP site (ftp://ftp.broad.mit.edu/pub/cmap/ratioMatrix.txt). The values of each element in the data matrix are fold-change ratios for a probe-set in a differential expression profile; we performed log_2_-transformation on these values. As each row in the data matrix is an Affymetrix GPL96 probe-set, we mapped the probe-sets to genes using the annotation file obtained from Gemma (https://gemma.msl.ubc.ca/arrays/showArrayDesign.html?id=1) (Zoubarev et al., 2012). In cases where multiple probe-sets map to a single gene, for each individual compound profile, we used the probe-set with the greatest magnitude of change in expression. Data for probe-sets that mapped to multiple genes were removed.

A profile is considered “unique” based on a combination of four experimental factors: the chemical compound used, dosage, treatment time and cell line used. An example would be “10μM metformin in MCF7 cells for 6 hours”. To avoid over-representation of a particular condition, in cases where there were duplicate profiles in the corpus (due to the same treatment-control condition being tested multiple times), a representative profile was selected at random. The resulting data matrix contains data for 12010 genes and 3742 profiles (representing 1309 compounds in up to 6 different doses in up to 5 different cell lines and 2 different durations). We refer to this as the CMap 1 data matrix.

### Processing of CMap 2 data

For CMap 2, the LINCS team provide two separate data releases known as Phase 1 and 2 in NCBI GEO (GSE92742 and GSE70138 respectively); we used both after merging them together. For Phase 2 data in particular, the version from “2017-03-06” was used. For gene-level metadata, the “Gene_Info” file is obtained from the GEO website. We used two versions of the CMap 2 differential expression data: first, the moderated z-score data provided by the LINCS team (in their parlance, “Level 5” data); second, we computed a log_2_-fold change data from the original “Level 3” expression data. The latter version was created for better comparability to CMap 1. Both are described in greater detail in the following paragraphs.

For the moderated z-score version of CMap 2, “Level 5” data was obtained from the GEO website. Metadata was obtained from the “Sig_Info” file associated with the GEO record. We filtered the data for perturbagen type “trt_cp” (i.e. “treatment with chemical compound”) and removed duplicate profiles in the corpus (as described for CMap 1 above). The resulting data matrix contains data for 12328 genes and 275335 profiles (representing 20547 compounds of 2750 different concentrations in 83 different cell lines and 5 different treatment durations, treated in various non-exhaustive combinations). We refer to this matrix as CMap 2-MZS (for “moderated z-score”).

To compute log_2_-fold changes, the “Level 3” data was obtained from GEO along with the associated metadata in the “Inst_Info” file. In LINCS parlance, “Level 3” data is more directly comparable to Affymetrix log_2_-transformed expression data. As for the CMap 2-MZS data, we selected samples with perturbagen type “trt_cp”, and also retained control samples with treatments “DMSO”, “PBS”, “H2O” and “UnTrt” (dimethyl sulfoxide, phosphate buffered saline, water and untreated, respectively). We then performed differential expression analysis as follows. For each treatment-control profile, we identified samples in which both the treatment and control samples are drawn from the same “RNA Plate Prefix” pool (e.g. “CPC001”, “CPC002”, etc.) Each treatment and control group were also restricted to have between 2 and 20 samples each, and to have equal number of samples between both treatment and controls; samples were drawn randomly when the number of samples exceeded these constraints. Differential expression for each gene was then computed by taking the difference of the mean expression between the treatment and control groups for each comparison. Similar to CMap 2-MZS, additional filtering was used to remove duplicate profiles in the corpus. The resulting data matrix contains data for 12328 genes and 275542 profiles (representing 20532 compounds of 3013 different concentrations in 83 different cell lines and 5 different treatment durations, treated in various non-exhaustive combinations). We refer to this matrix as CMap 2-FC (for “fold change”).

### Harmonization of CMap 1 and 2 data

The goal of harmonization was to identify genes and conditions that were common across the datasets. Conditions were considered the same if they involve application of the same compound at the same concentration to the same cell line for the same duration of time, leading to a list of 109 conditions used for downstream analysis (Table S1). Next, we identified genes that were common between CMap versions and were also labelled as “BING” genes in the CMap 2 metadata (“BING”: Best Inferred Genes, which includes landmark genes that were directly measured, and genes whose expression was considered to be predicted with high accuracy (Subramanian et al., 2017)). This resulted in our harmonized datasets having differential expression information for 11,689 genes, 963 of which are landmark genes.

### Querying CMap 2 with CMap 1 signatures

The CMap 2 corpus was queried through the L1000-Query web interface, a tool provided by the LINCS team for registered users. The corpus included in the web tool contains a subset of Phase 1 profiles derived from 8,388 tested perturbagens (2,429 of which are compounds), known as the “Touchstone” subset. We chose the CMap 1 signatures for querying against the CMap 2 corpus through a number of filtering steps. Starting from the initial 3,742 unique CMap 1 profiles, we selected representative profiles for each combination of chemical compound and cell line by picking the profile with the highest dosage. Next, we constrained the cell line to either be for MCF7 or PC3, the treatment time for 6 hours, and the compounds that are common between CMap 1 and the CMap 2 “Touchstone” subset; this is to reflect the conditions that are available in the CMap 2 corpus used in L1000-Query.

The L1000-Query interface has a few requirements: it requires two gene lists (for over- and under-expression) as input, and it only recognizes “BING” genes. It also has a gene list size restriction of between 10 and 150 genes. Thus, we filtered our CMap 1 data to exclude non-BING genes, and used a log_2_-fold change threshold of 1.0 and −1.0 to identify over- and under-expressed signatures respectively. Conditions which had too few or too many differentially expressed genes were excluded. This gave us a shortlist of 588 signatures (Table S2). Each was then used to query L1000-Query and the compound ranking results (in the form of signature similarity scores or “Tau scores” in LINCS parlance) were exported. We then filtered the ranking results for compounds that are present in CMap 1.

Separately, we performed the same analysis, but using CMap 1 signatures derived from the harmonized data instead, of which 99 of the 109 conditions were represented in the L1000-Query “Touchstone” subset (Table S3). Additionally, we used four different thresholding procedures to determine the genes included in the query signatures: “Hybrid Threshold”, “Top-100”, “Top-20” and “Top-300”. The “Hybrid Threshold” method first uses a log_2_-fold change threshold of 1.0 and −1.0 to identify over- and under-expressed genes. If the threshold resulted in gene hit lists (over-/under-expressed lists separately) containing more than 150 genes, the genes are then ranked by magnitude of change (over-/under-expressed separately) and the top-150 genes are selected instead. In the converse situation, where either gene hit lists contain less than 10 genes, the condition is not used to query CMap 2; two conditions (out of 99) were disqualified by this criterion: “10μM rosiglitazone in MCF7” and “10μM scopolamine in PC3”. For the “Top-100” method, unlike all other thresholding methods which uses all genes, uses only the landmark genes; the landmark genes are first ranked by magnitude of change, and the top-50 over-expressed and top-50 under-expressed genes are selected (as such, a total of 100 genes are used in the query). This thresholding method is also used by the LINCS team for pre-calculating the connectivity between compounds included in the Touchstone subset (Subramanian et al., 2017). The “Top-20” and “Top-300” thresholding methods are similar to that of “Top-100”, except that all genes (instead of just landmark genes) are used, and either the top-10 or top-150 genes (for over- and under-expressed gene lists separately, giving a sum of either 20 or 300 genes) are selected for each method respectively.

### Literature-derived signatures

We searched the literature to identify reports where CMap 1 had been applied to a novel gene signature, and in which the results were validated with an independent assay. We omitted papers which do not provide the actual gene signature, or which had too few genes to be eligible for the L1000-Query tool. In the end we identified usable signatures from two studies (Braconi et al., 2009; Brum et al., 2015). Each was then used to query L1000-Query and the results filtered for compounds present in CMap 1 to enable fair comparison. The ranks were then normalized as described above.

Similarly, we also searched literature to identify reports where CMap 2 had been applied to a novel gene signature, and in which the results were validated with an independent assay. We omitted papers which do not provide the actual gene signature, or which the validated drugs were not included in CMap 1. We identified usable signatures from one study (Ferguson et al., 2018); the signatures were then used to query CMap 1 as described below.

The CMap 1 corpus was queried through the old CMap portal’s (https://portals.broadinstitute.org/cmap/) web interface provided for registered users. Unlike CMap 2, CMap 1 does not impose any limitations on the number of Affymetrix GeneChip probe-sets used in the query. We queried CMap 1 with the 16 signatures used in Ferguson et al., and exported the results. The results were not constrained by cell line and were combined using the “75^th^ percentile ordering” and “rank-product” methods, as described by Ferguson et al. (Ferguson et al., 2018).

### Modified-Jaccard index for measuring similarity

While Spearman rank correlation (r_s_) is used as the main measure of similarity between data vectors throughout this paper, we report a modified Jaccard index as an alternative measure. The classical Jaccard index (or Jaccard similarity coefficient) is defined as the ratio of the number of intersecting elements between sets A and B over the number of shared (or union) elements between sets A and B. This is mathematically expressed as:

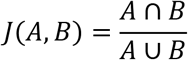

Where 0 ⩽ J(A, B) ⩽ 1, and if A ∪ B is empty, J(A, B) = 1. Alternatively, we could use A and B to represent vectors of binary values (of length *n*), each element (indexed *i*) in the vectors indicating presence or absence (values ∈ {0, 1}); coupled with the Boolean operators “AND” (∧) and “OR” (∨), this gives us the following expression:

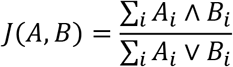

As the Jaccard index operates on Boolean or binary values (i.e. absence/presence), the measure as-is is non-ideal for our thresholded differential expression data, which is expressed in ternary data values, i.e. −1, 0 and 1 for under-expressed, unchanged and over-expressed respectively.

Thus, we modified the Jaccard index to accommodate ternary data values, and it is represented below:

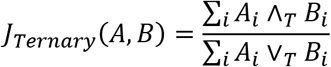

Where ∧_T_ is the ternary “AND” operator and ∨_T_ is the ternary “OR” operator. For the ternary “AND” operator, it returns 1 if both input values (i.e. A_i_ and B_i_) are concurrently either −1 or 1 (i.e. both genes are differentially expressed AND changes in the same direction), otherwise it returns 0. The ternary “OR” operator returns 1 if either input value is −1 or 1, otherwise it returns 0 if both inputs are 0 (i.e. if either gene is differentially expressed, it is included in the count). Additionally, instead of returning a Jaccard index of 1 if both sets A and B are empty, our modified Jaccard index function would return 0. This is based on the principle that the modified Jaccard index is supposed to capture the agreement of differentially expressed gene lists between two vectors: if no genes are differentially expressed in both lists, it is not meaningful in our use case to return a value 1, since it means that “both gene lists are similarly not-differentially expressed”, and would be indistinguishable from the actual case of interest, which is “both gene lists have exactly the same differentially expressed genes”.

In order to apply the modified-Jaccard function on the differential expression data, the data itself needs to be thresholded to generate the ternary values. The thresholds used for CMap 1 and CMap 2-FC data is −1.0 and 1.0 for under-expressed and over-expressed respectively. CMap 2-MZS data is first “de-standardized” by multiplying the z-scores with the square root of the number of contributing sample replicates, followed by a threshold of −2.0 and 2.0 for under-expressed and over-expressed respectively, as described in the original publication (Subramanian et al., 2017).

### Within-dataset correlation of differential expression profiles

The CMap 1 and 2 differential expression data used for this analysis was processed as outlined above (see “Processing of CMap 1/CMap 2 data”), with the exception that duplicate conditions were retained. We then filtered the data for cell lines “MCF7” or “PC3”, and treatment time of “6 hours”. Conditions where there is only a single profile instance (i.e. no replicates) were removed. The resulting number of unique replicated conditions for CMap 1, CMap 2-FC and CMap 2-MZS are 1614, 2416 and 2477 respectively. Pairwise similarity (Spearman correlation and modified-Jaccard index) of the profiles, by condition, was then performed; a total of 7809, 9020 and 10630 comparisons were generated for CMap 1, CMap 2-FC and CMap 2-MZS respectively. We performed multiple downstream analyses using this data.

First, we investigated the distribution of pairwise correlation of profiles. In this analysis, we took the maximal correlation by condition as the summary statistic to correct for unequal numbers of replicate comparisons for each condition. In related analyses where we establish the relationship between CMap 2 compound retrieval performance to *cross*-dataset agreement of differential expression to *within*-dataset agreement of differential expression, the same summary statistic (i.e. maximal correlation) is also used.

Next, we assessed the relationship between within-dataset agreement of differential expression and the number of differentially expressed genes in each differential expression profile. The threshold for determining if a gene is differentially expressed is outlined above (see “Modified-Jaccard index for measuring similarity”). Since each profile replicate comparison consists of a pair of differential expression profiles, and each profile will have its own number of differentially expressed genes, we chose the lesser number of differentially expressed genes as the representative value for each comparison-pair. The rationale is that the profile with less differentially expressed genes would have a weaker “differential expression strength”, and as such reducing the likelihood of the pair having high profile agreement (i.e. acting as the “limiting profile”).

Additionally, we also evaluated the relationship between the pairwise correlations and compound concentration. In this analysis, we restricted our analysis to correlations for conditions that have at least three profile replicates and compounds with at least three different concentrations. For CMap 1, only one compound (i.e. valproic acid) tested in the MCF7 cell line was suitable; For CMap 2-MZS, 10 and 7 compounds (tested in the MCF7 and PC3 cell lines respectively) were identified in the initial screen – we further restricted the analysis to the four compounds that had adequate data in both MCF7 and PC3 cell lines (i.e. vorinostat, trichostatin A, geldanamycin and wortmannin).

### Within-dataset correlation of sample-level gene expression profiles

As we used CMap 1’s pre-computed differential expression data directly instead of the raw expression data, this analysis is limited to CMap 2 only. Additionally, calculation of within-dataset correlation of sample expression profiles is limited to the “differential expression profile pairs” used in the calculation of within-dataset correlation of differential expression described above. Specifically, since the replicate-pairs consist of two differential expression profile replicates of the same condition, and each differential expression profile in the comparison-pair is “constituted” from a group of samples (“treatment” sample only in the case of CMap 2-MZS; “treatment” and “control” samples for CMap 2-FC), the “agreement of sample expression” is calculated using Spearman correlation, but represented in two manners. In the first case (henceforth referred to as “in-replicate”), for each differential expression replicate, the maximum correlation of gene expression between its constituent samples is calculated; the representative sample correlation for the differential expression replicate-pair would be the smaller value between the two maximums (i.e. “limiting correlation”). The rationale is that the replicate with the lower maximal sample correlation would impose an upper threshold on the agreement between two differential expression replicates (i.e. a “noisy” profile is unlikely to agree with a non-“noisy” profile). In the second case (henceforth referred to as “cross-replicate”), the sample correlation is calculated by comparing the gene expression of the constituents of one differential expression replicate to that of the other replicate (in the comparison-pair) in a pairwise manner, then taking the maximum correlation of the result. The idea is that the maximum (“best case”) cross-replicate agreement of sample gene expression would be a proxy to the agreement of the differential expression replicate pairs themselves. As the sample expression profile agreement is calculated based on the differential expression replicate pairs outlined earlier, the total number of sample agreements are 9020 and 10630 for CMap 2-FC and CMap 2-MZS respectively; the number of sample agreements by unique conditions are 2416 and 2477 for CMap 2-FC and CMap 2-MZS respectively.

We used a multivariate linear regression to model the relationship between within-dataset agreement of differential expression and three different predictors: the number of differentially expressed genes, the “in-replicate” and “cross-replicate” sample expression agreement. The linear model we used is expressed below:

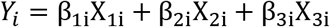

Where for each unique condition *i* (out of the 2477 unique conditions in CMap 2-MZS with within-dataset reproducibility results), *Y*_*i*_ is the within-dataset agreement of differential expression; *X*_*1i*_ is the number of differentially expressed genes, *X*_*2i*_ is the “in-replicate” agreement of sample expression, *X*_*3i*_ is the “cross-replicate” agreement of sample expression; β_1i_, β_2i_ and β_3i_ are the respective parameter coefficients. We report the standardized regression coefficients (Β) that were calculated using the *lm.beta* R package.

### Processing of the DeAbrew dataset

The data for the GEO record corresponding to De Abrew et al. (De Abrew et al., 2016), GSE69845, was loaded into the Gemma platform (http://tinyurl.com/Gemma-GSE69845). The raw data was reprocessed from CEL files using the “RMA” algorithm and differential expression analysis was performed for each treatment versus control using a linear model. The computed log_2_-fold change values were used for downstream analysis. For comparing to the CMap 1 and 2 data sets, we manually identified 12 of the 34 compounds used by De Abrew et al. in which the concentrations (10μM) are the same or very close (10 ± 5μM, i.e. difference < 50%) to their counterparts in CMap 1 and 2 (Table S4).

## Results

### CMap 2 does not reproduce CMap 1 drug prioritization

We first asked whether CMap 1 and 2 give concordant results in their core use case: drug prioritization. We first did this by querying CMap 2 through LINCS’s L1000-Query website, with hit lists derived from CMap 1 profiles. We reason that if the two datasets were concordant, given a signature for a compound from CMap 1, when used to query CMap 2, should yield that same compound as a top result. In fact, this analysis represents a best-case scenario for CMap 2. If CMap 2 performs well at this task, there would be increased confidence in real-life applications that use queries from more heterologous contexts.

In this analysis, we considered compounds that were present in both CMap 1 and 2. To select the CMap 1 signatures, we chose the highest compound concentrations available, yielding 588 signatures (see Methods and Table S2). These were used as inputs to query against CMap 2. We also performed the control experiment, querying L1000 with profiles derived from CMap 2 for the same 588 conditions; as the query results were pre-computed by LINCS and provided online as the Touchstone database, we just retrieved the pre-computed results. We predict that these “self-queries” should highly rank the compound matching the condition used in the query, and the results representing the upper-bound of CMap 2’s retrieval performance.

The results are shown in Figure 1. In our control comparison, querying CMap 2 with a profile taken from CMap 2 data results in the correct compound being prioritized in the top-10% (i.e. rank ≤ 68) 83% of the time, or 486 out of 588 profiles (blue lines in Figure 1). Using a stricter threshold more reflective of real-life applications, 53% or 313 profiles have the correct compound ranked first. These results provide a benchmark for the primary test, which was to probe CMap 2 with signatures from CMap 1 (red lines in Figure 1). In this test, the correct compound was ranked in the top-10% for only 99/588 signatures (17%), with only five compounds ranked first (<1% of signatures; these five were 15-Δ-prostaglandin J2, flumetasone, geldanamycin, niclosamide and sirolimus).

**Figure 1:**
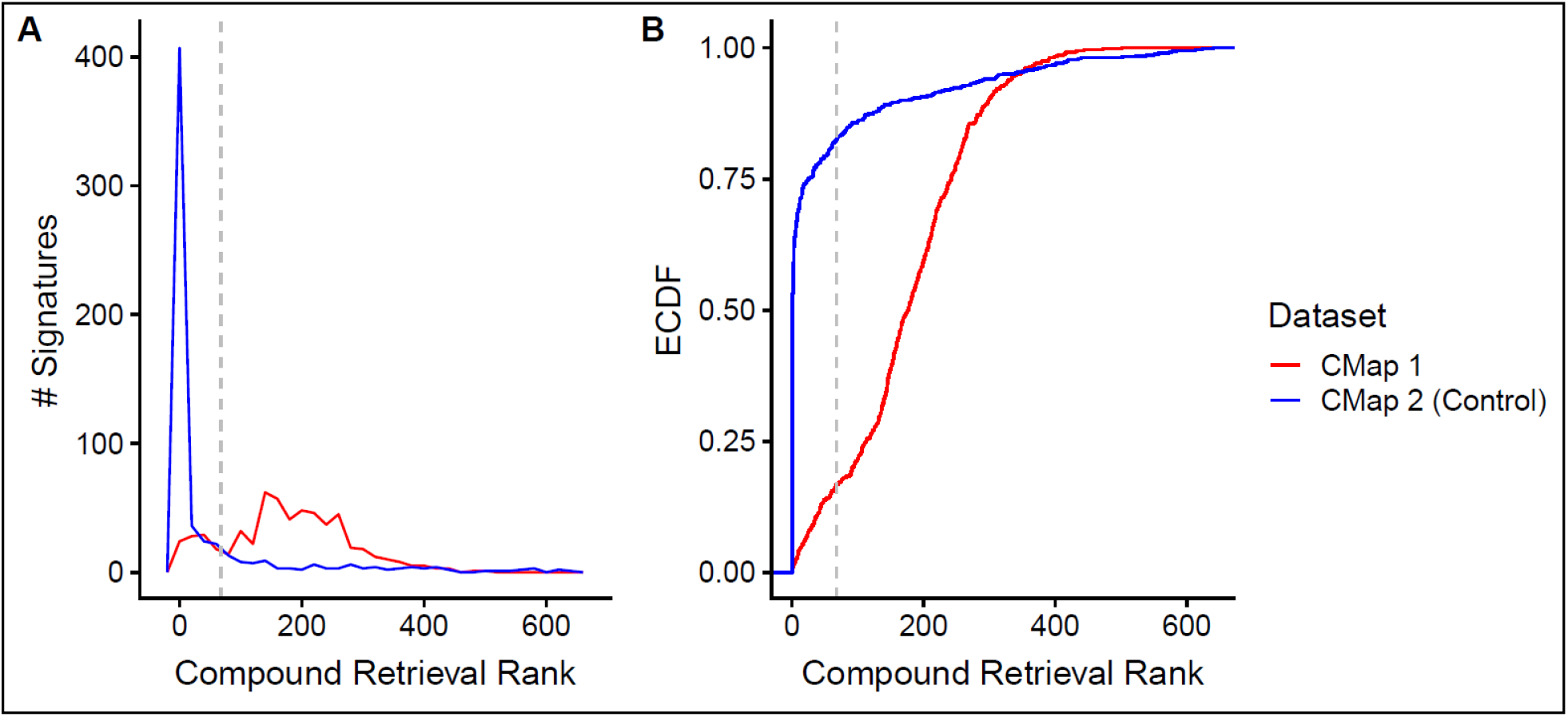
Distribution of compound retrieval ranks (1 is best, x-axis) from querying CMap 2 using signatures derived from CMap 1 (red lines) or CMap 2 (blue lines; self-query) data, (N = 588). A shows the distribution while B is the empirical cumulative distribution function (ECDF) of the same results. The dotted grey line indicates the profiles where the query compound is ranked in the top 10% (rank ⩽ 68).

We then examined the data for factors that might explain differences in performance for different queries. CMap 2’s retrieval performance using CMap 1 signatures is modestly predicted by the number of DE genes (rank correlation, r_s_ = −0.24; p = 5.3 × 10^−9^; Figure S3A). There was also a small effect of cell line, with profiles derived from the PC3 cell line (median rank, r_med_ = 159) performing slightly but significantly better than their MCF7 counterparts (r_med_ = 188; Mann-Whitney U Test p-value = 5.6 × 10^−3^); the same is also observed in the self-query controls (r_med_ for PC3 cell line = 1, MCF7 cell line = 2; p = 2.2 × 10^−6^). These results suggest that DE strength (i.e. number of DE genes) and cell line-specific effects may affect CMap 2 retrieval performance when using CMap 1 signatures.

In the above analysis, we did not require that the same compound concentration used in CMap 1 be available in CMap 2. Thus, we repeated the analysis, but using CMap 1 signatures derived from the harmonized data (see Methods and Table S3) which ensured that the compound signatures of a particular concentration is present in both CMaps. Additionally, we tested multiple thresholds for generating the query signatures to assess the effect of signature sizes on retrieval performance. From the harmonized data, a total of 99 signatures were generated (with the exception of “Hybrid Threshold”, which had two less signatures due to an inadequate number of DE genes). Regardless of thresholds used, the initial findings were reproduced. Using the query results of CMap 1 “Hybrid Threshold” signatures as an example, the correct compound was ranked in the top-10% for 37 signatures (38%); and ranked first for 12 signatures (12%). In comparison to the CMap 2 self-query, the correct compound was ranked in the top-10% for 82 signatures (85%); and ranked first for 54 signatures (56%; Figure S1). The number of DE genes also predicts CMap 2’s retrieval performance using CMap 1 signatures in this subset (r_s_ = −0.26, p =1.2 × 10^−2^; Figure S3B).

Based on these results, it is apparent that CMap 2 is usually unable to accurately prioritize compounds based on CMap 1 signatures for the exact same compound under the same conditions, which should be a best-case scenario for CMap 2.

We also assessed if CMap 2 retrieval performance is affected by query signature sizes. In Figure S2, we see that thresholds resulting in greater CMap 1 signature sizes tend to have better retrieval performance: “Top-300” and “Top-100”-derived signatures have the correct compound ranked in the top-10% for 39 and 37 signatures respectively (out of 99); while “Top-20”-derived signatures in only 26 signatures. The median retrieval rank as an alternative metric shows similar trends: 110 and 144 for “Top-300” and “Top-100” respectively, 241 for “Top-20” (r_med_ of “Hybrid Threshold” = 108). This confirms that querying CMap 2 with larger signatures increases reproducibility of drug recommendations.

We next considered if there is any relationship between CMap 2 self-retrieval and retrieval of CMap 1 signature queries. While the correlation between the two are rather weak (r_s_ in 588-conditions data = 0.17; “Hybrid Threshold” 97-conditions = 0.20), we see a trend where conditions that have limited CMap 2 self-retrieval performance (rank > 68) also have limited retrieval performance when queried with CMap 1 signatures (Figure S4). The converse does not hold true however: conditions with good CMap 2 self-retrieval performance do not guarantee equally good retrieval performance when queried with CMap 1 signatures. These observations suggest that CMap 2 users can further refine compound recommendations by assigning lower confidence to compounds that have limited self-retrieval performance.

Our second test involved querying CMap 2 using hit lists derived from published applications of CMap 1. We identified studies in which the authors queried CMap 1 with a signature they generated from other data, followed by experimental verification of the biological effects of the recommended compounds relevant to their study. We were able to identify two suitable signatures (Braconi et al., 2009; Brum et al., 2015). We hypothesized that if CMap 2 was performing well, it should highly rank the same compounds as identified and validated in the publications.

In Brum et al., CMap 1 was queried with a signature derived from human mesenchymal stromal cell-derived osteoblasts (Brum et al., 2015, 2018). Among the top compounds recommended were parbendazole, withaferin-A and amylocaine. We used L1000-Query to test whether their input signature would retrieve these compounds from CMap 2. As the hits from CMap 1 were associated with PC3 cells, we constrained the results to that cell line, and we only ranked the 628 compounds shared in both CMap 1 and 2. These constraints were intended to make the comparison as fair as possible. In Table 1, we see that parbendazole, the key finding of the 2015 paper, was ranked 142 by CMap 2 (Tau Score, τ = 16.12; τ being the normalized measure of signature similarity provided by the CMap software, with scores > 90.0 being highly similar and < −90.0 being highly dissimilar: thresholds as recommended by (Subramanian et al., 2017)). Withaferin-A and amylocaine were also not highly ranked by CMap 2 (ranks of 196 and 122 respectively; τ = 0 and 22.86 respectively, Table 1). In contrast, known osteogenic glucocorticoids were ranked in the top-20 in both CMaps.

**Table 1:**
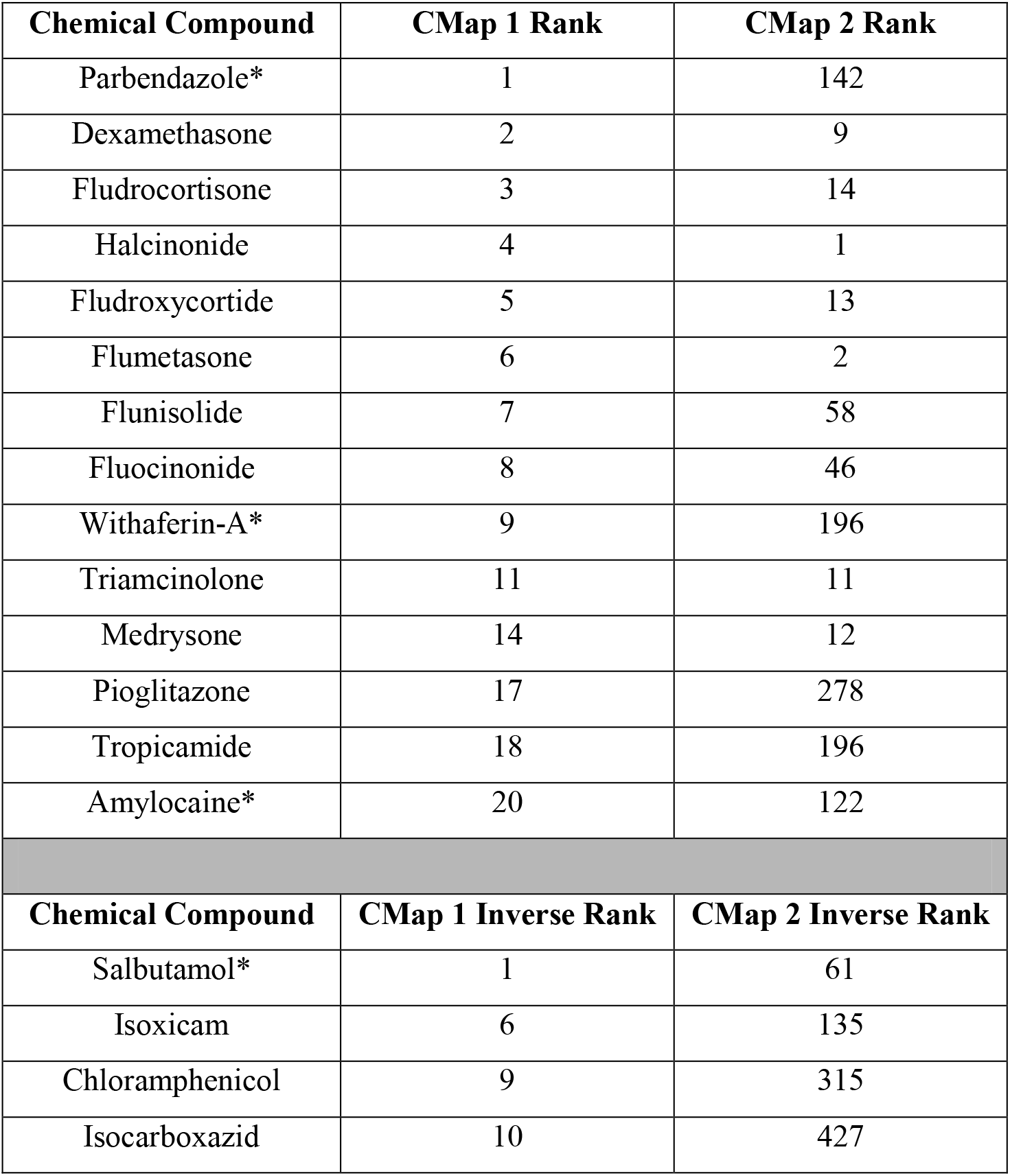
Compound rankings from GSEA-analysis using previously reported signatures from Brum et al. (2015 and 2018). Inverse-ranks are reported for “reversing” the query signature. Compounds marked with asterisks (*) were experimentally validated. The CMap 2 ranks are relative to CMap 1 compounds contained in CMap 2.

Brum et al. (2018) were also interested in compounds that could reverse the query signature and validated the dexamethasone-inhibiting activity of salbutamol, which was recommended by CMap 1; CMap 2 failed to recommend this compound, giving an inverse-rank of 61 (τ = −68.41).

We did a similar analysis based on the report of Braconi et al. (Braconi et al., 2009). In that study, 17-allylamino-geldanamycin (17-AAG) and resveratrol were recommended by CMap 1 in response to a query designed to identify inhibitors of vascular invasion by hepatocellular cancer. For their analysis, Braconi et al. were looking for drugs that yielded signatures that “reversed” the query signature, so compounds with negative “Connectivity Scores” were of interest (“Connectivity Scores” in CMap 1 is analogous to CMap 2’s τ, except that the range is different: “−1 to 1” for CMap 1, “−100 to 100” for CMap 2; and CMap 2’s τ have been corrected for non-specificity of signature similarity). While the cell line context was not reported by the authors (and there are no liver cancer cell lines in CMap 1), as the bulk of their subsequent biological assays were performed on HepG2 cell lines, we constrained our L1000-Query rankings for that cell line. As before, we only considered compounds shared between both CMaps (N = 1904). Unfortunately, 17-AAG was not included in CMap 2’s Touchstone subset, so the reproducibility of that finding could not be directly assessed. As 17-AAG is a geldanamycin-derivative, we included geldanamycin as a “stand-in” in the CMap 2 analysis. In Table 2, we see that both resveratrol and geldanamycin have a negative τ, which is consistent with negative connectivity scores reported by Braconi et al. However, resveratrol was ranked 419, which is inconsistent with the CMap 1 results reported by Braconi et al. The results for geldanamycin was more encouraging, with a rank of 18, but it is notable that geldanamycin was not identified by Braconi et al. when querying CMap 1, despite its inclusion in CMap 1.

**Table 2:**
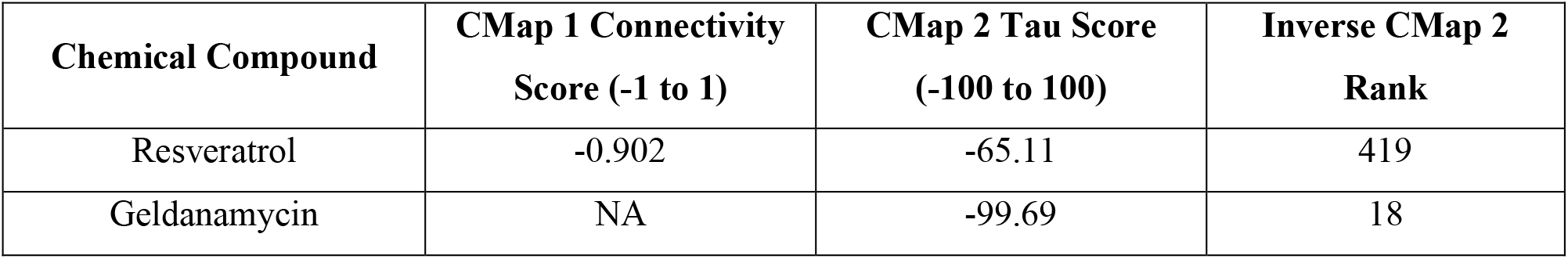
Compound rankings from GSEA-analysis using previously reported signature from Braconi et al. (2009). Inverse-ranks are reported for “reversing” the query signature. As with Table 1, CMap 2 ranks are relative to CMap 1 compounds contained in CMap 2.

| Chemical Compound | CMap 1 Connectivity<br>Score (-1 to 1) | CMap 2 Tau Score<br>(-100 to 100) | Inverse CMap 2<br>Rank |
| --- | --- | --- | --- |
| Resveratrol | -0.902 | -65.11 | 419 |
| Geldanamycin | NA | -99.69 | 18 |

We next reversed the test by querying CMap 1 using signatures derived from a published application of CMap 2 (Ferguson et al., 2018). Similar to the original test, we hypothesized that if CMap 1 is performing similarly well as CMap 2, it should highly rank the same compounds as identified and validated in the publication.

In Ferguson et al., CMap 2 was queried with signatures of brains from a mouse model of alcohol use disorder (HDID-1 strain) to identify compounds that can induce reduction in alcohol intake (Ferguson et al., 2018); pergolide was recommended and subsequently experimentally validated. Similar to the Braconi et al. study, the authors were looking for compounds that would reverse the query signature, so compounds with negative “Connectivity Scores” were of interest. Unlike the previous cases we examined, the query signatures are derived from mouse brain tissue, not human cell lines, thus we did not constrained our CMap 1 rankings for cell line specificity. We however still only considered compounds shared between both CMaps (N = 663). Additionally, Ferguson et al. used three different meta-analytical algorithms separately to collapse the compound rankings into one hybrid ranking; we used only two of them (i.e. “75^th^ percentile” and “rank-product”) as the third method was not applicable to CMap 1. In Table 3, we see that CMap 1 failed to rank highly the primary finding, pergolide. The same is observed for the two other drugs, with the exception of alvespimycin when using the “75^th^ percentile” meta-ranking, with a ranking of 15. As the authors were reporting the top-15 recommended drugs for each summarization method, this is the only successful observation of a reproducible result.

**Table 3:** Compound rankings from GSEA-analysis using previously reported signatures from Ferguson et al. (2018) that is further summarized using the “75^th^ Percentile” and “Rank-Product” methods as described in the original publication. Inverse-ranks are reported for “reversing” the query signatures. Compounds marked with asterisks (*) were experimentally validated. The CMap 1 ranks are relative to CMap 2 compounds contained in CMap 1.

| Chemical Compound | “75<sup>th</sup> Percentile” Inverse Rank |  | “Rank-Product” Inverse Rank |  |
| --- | --- | --- | --- | --- |
|  | CMap 2 | CMap 1 | CMap 2 | CMap 1 |
| Pergolide* | 1 | 269 | 3 | 138 |
| Alvespimycin | 5 | 15 | 5 | 102 |
| SB-203580 | - | 135 | 4 | 145 |

Taken together, these results suggest that the consistency between prioritization results from CMap 1 and CMap 2 is low. The most likely source of this discordance is the underlying data, which we investigated next.

### Low cross-dataset DE agreement is predictive of limited CMap 2 retrieval performance

To assess the similarity of data for conditions shared between both CMap 1 and 2 (N = 109, Table S1), we computed the rank correlations of the DE profiles between CMap 1 and 2 for each condition. We considered if the two versions of the CMap 2 data (fold-change in CMap 2-FC and z-scores in CMap 2-MZS, see Methods), as well as whether limiting the data to the 963 “landmark genes” directly measured in CMap 2 had any impact. The results are shown in Figure 2. Regardless of how CMap 2 data was processed or the genes used, the mean CMap 1 – CMap 2 correlation ranges from 0.05 to 0.11 (Figure 2), with only a few exceptions (2 and 7 comparisons with r_s_ > 0.6 using landmark genes; Figure 2C and 2D respectively). The three compounds that had the highest agreements (r_s_ > 0.6) were vorinostat, trichostatin-A and scriptaid. Similar results are observed even when limiting the conditions to compounds that are only available in the Touchstone subset (Figure S5). Scatter plots of the data for representative pairs of profiles with low, medium and high agreement are shown in Figure 3. Generally, cross-dataset DE reproducibility between CMaps is low.

**Figure 2:**
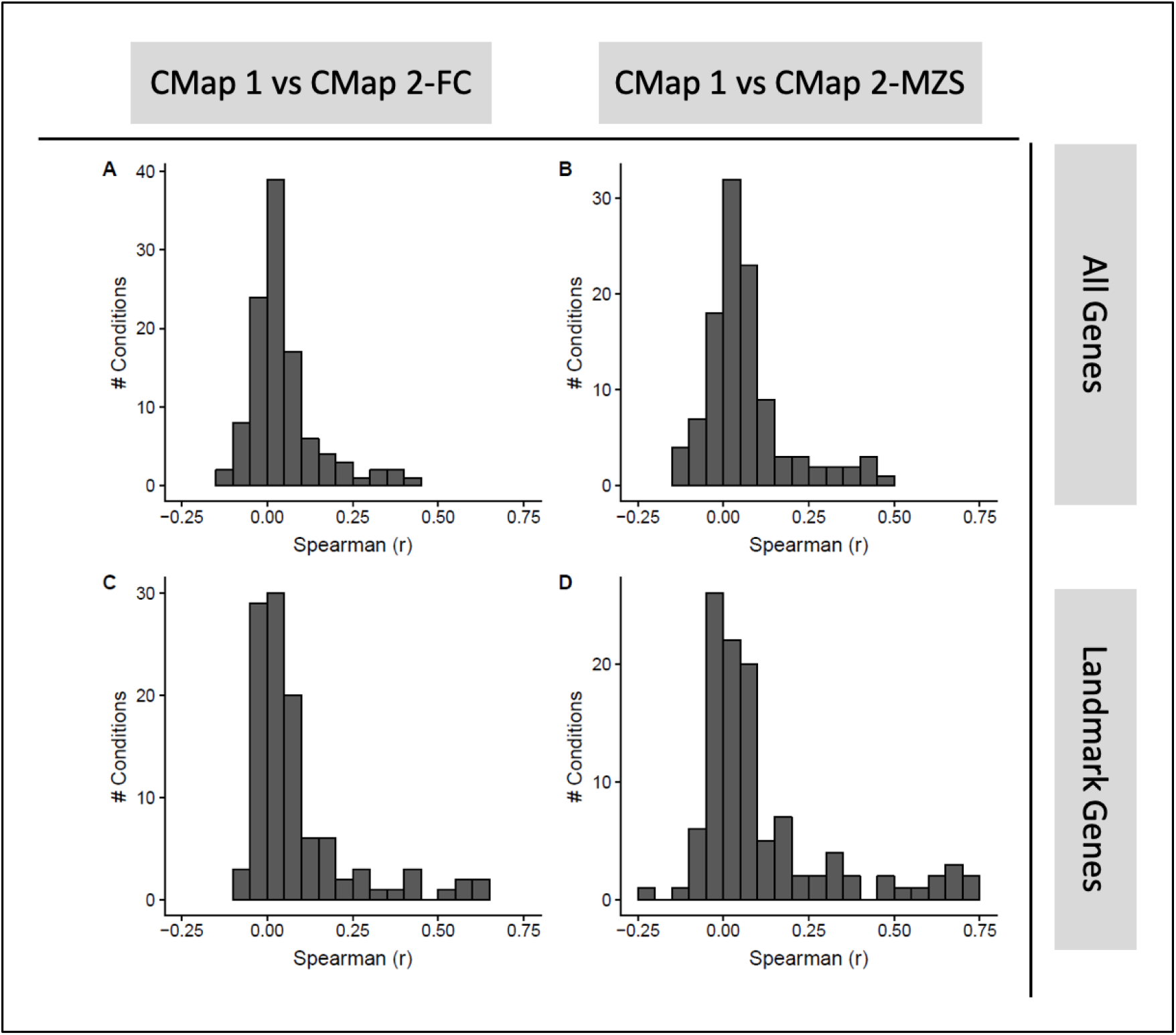
Distribution of pairwise rank correlation of DE profiles from CMap 1 against CMap 2. A and C are comparisons against CMap 2-FC; B and D against CMap 2-MZS. All genes common between CMap 1 and 2 are used in A and B, while only “landmark genes” are used in C and D. (N = 109 conditions).

**Figure 3:**
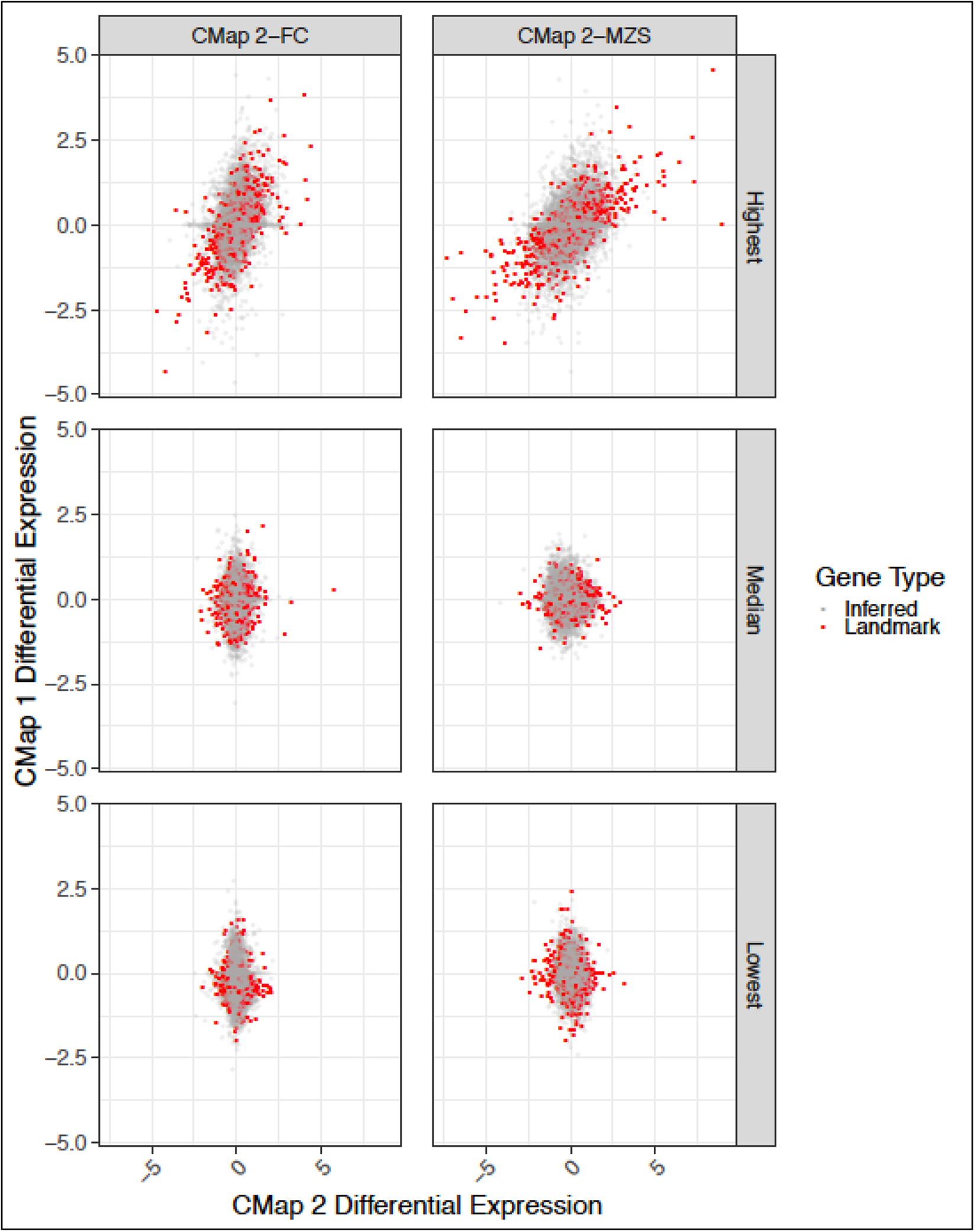
Representative scatter-plots of gene DE values for six different pairs of profiles with different levels of between-CMap agreement (CMap 1 – CMap 2; high, medium or low). Each point represents one gene. The x-axis are values derived from CMap 2 profiles and y-axis are from CMap 1. The plots from top-to-bottom are of profiles with the highest, median and lowest pairwise rank correlation between CMaps respectively; plots on the left use the fold change version of CMap 2 (CMap 2-FC) while those on the right use the moderated Z-score version (CMap 2-MZS). Genes where the expression level is measured in CMap 2 (“Landmark genes”) are coloured red while those that are inferred are in grey. In order from top-to-bottom and from left-to-right, the profiles are (values in bracket are rank correlation of landmark genes): 1μM trichostatin-A in MCF7 (r = 0.63), 10μM vorinostat in MCF7 (r = 0.72), 10μM 16,16-dimethylprostaglandin-E2 in PC3 (r = 0.03), 10μM SR-95639A in PC3 (r = 0.04), 10μM orlistat in PC3 (r = −0.10) and 10μM resveratrol in MCF7 (r = −0.22). Treatment time for all profiles was 6 hours.

One concern raised by Figure 3 is that many genes in the profiles have very little DE (i.e. values close to zero) and that rank correlation may be a suboptimal assessment metric (due to a majority of ranks being “noisy”). We used rank correlation primarily because the underlying analytical approach in the CMap software is based on gene ranks, rather than gene list overlaps that requires an arbitrary threshold (Subramanian et al., 2005). Still, to ensure we were not underestimating agreement, we repeated the assessment using gene overlaps at an arbitrary threshold, for “all genes” and “landmark genes only” respectively (see Methods). As shown in Figure S6, the distribution of cross-dataset agreement is very similar to that in Figure 2, indicating that regardless of the similarity metric used, the overall reproducibility of DE profiles between both CMaps across the 109 shared conditions is low.

Next, we considered if poor cross-dataset agreement may be the cause for poor retrieval performance of CMap 2 when queried with CMap 1 signatures (Figure S1). Out of the 109 profiles, 97 CMap 1 profiles (“Hybrid Threshold” signatures, see Methods) were used to query CMap 2. We found that CMap 2 retrieval performance of CMap 1 signatures was correlated with cross-dataset agreement (r_s_ = −0.55, p =4.3 × 10^−9^; Figure S7A); there was however no relationship between CMap 2 self-retrieval performance and cross-dataset agreement (r_s_ = −0.004; Figure S7B). We also observed that once the cross-dataset agreement is above 0.2, CMap 2 is able to rank the correct compound in the top-10%, regardless which CMap the signature originated from. This shows that cross-dataset DE agreement is predictive of CMap 2 compound retrieval performance.

### Low within-dataset DE agreement in both CMaps is predictive of limited CMap 2 retrieval performance

The lack of replicability of the DE profiles between CMaps raised further questions about the quality of the data; specifically, the consistency of the DE profile replicates *within* the datasets. Thus, we investigated the similarity of the DE profiles for the same exact condition within each CMap version. In Figure 4, we see that regardless of version, the distribution of the maximum (“best case”) rank correlations for DE replicates is centered near zero, indicating most conditions in both CMaps have low within-dataset reproducibility (all mean r_s_ < 0.14); this level of agreement is lower than expected for cross-laboratory studies, such as the minimum (“worst case”) rank correlations (r_s_ = 0.90) observed in the MAQC formal evaluation of DE across laboratories (Shi et al., 2006). The agreement remains low even when using just landmark genes (all mean r_s_ < 0.18) or the modified Jaccard index (all mean J_T_ < 0.13, Figure S8). This shows that there is low within-dataset reproducibility.

**Figure 4:**
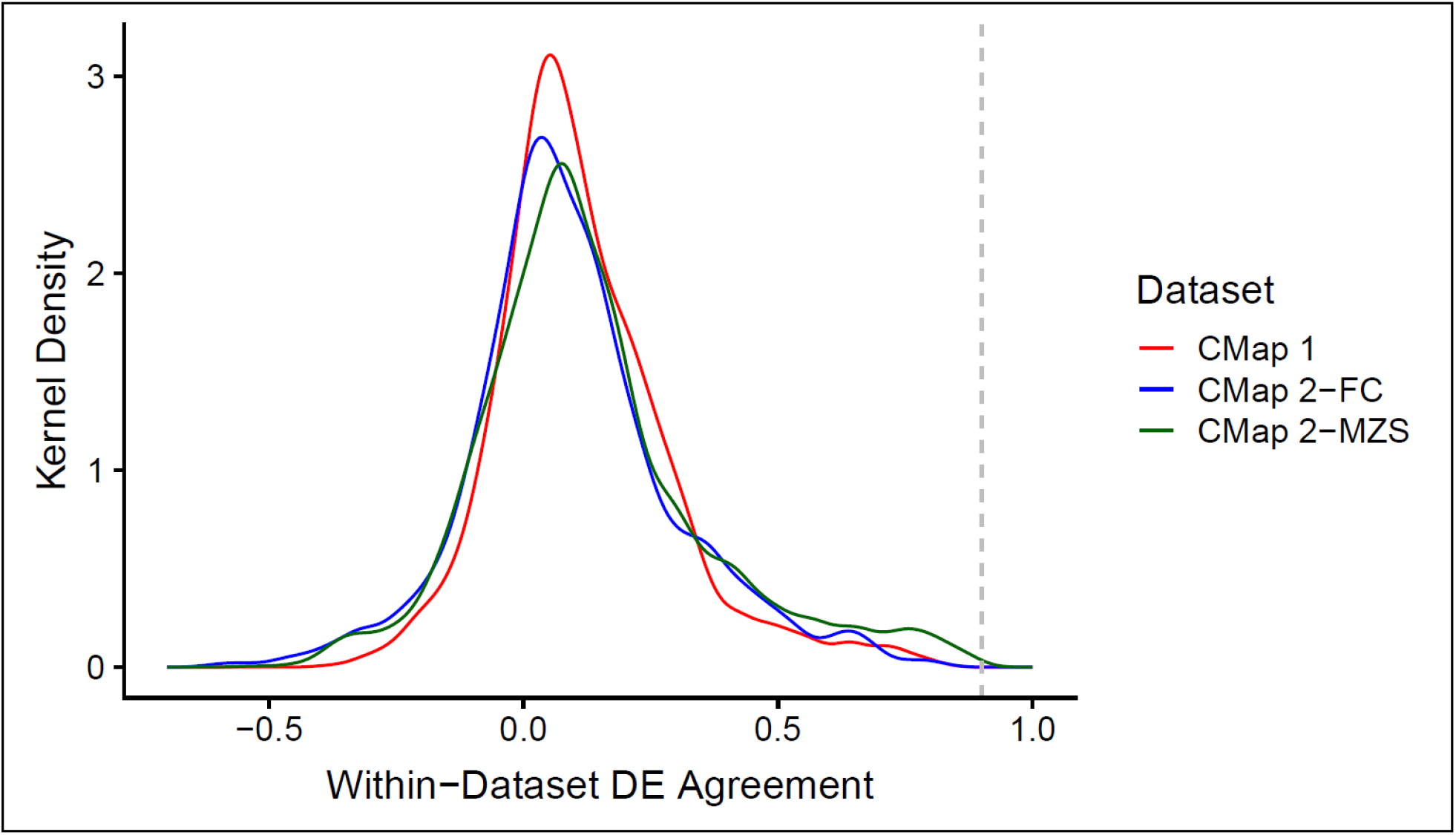
Distribution of maximum (“best case”) pairwise rank correlation for DE profile replicates of the same condition calculated using all genes, within each dataset. The number of unique conditions assessed for CMap 1, CMap 2-FC and CMap 2-MZS are 1614, 2416 and 2477 respectively. The dotted grey line indicates the minimum (“worst case”) rank correlation of DE for intra-platform cross-laboratory comparisons reported by MAQC (r_s_ = 0.90).

Seeking an explanation, we suspected that there is minimal DE occurring in many of the profiles themselves to begin with, thus causing the observed poor reproducibility: we found this to be true. For both CMap 1 and 2, the number of DE genes is predictive of a condition’s within-dataset agreement (r_s_ for CMap 1 = 0.40; CMap 2 = 0.32; Figure S9). This suggests that high DE reproducibility is contingent on having many DE genes.

Next, we considered if the poor within-dataset reproducibility is associated with CMap 2’s limited retrieval performance of CMap 1-signatures (Figure S1). Out of the 97 queried signatures, 65 and 58 signatures (for CMap 1 and CMap 2-MZS respectively) had adequate number of DE profile replicates in the respective CMap datasets for this assessment; we used all genes in the calculation. In Figure S10, we see that CMap 2’s retrieval performance of CMap 1 signatures is associated with within-dataset DE reproducibility in both CMap 1 (r_s_ = −0.56, p = 1.6 × 10^−6^, N = 65; Figure S10A) and CMap 2-MZS (r_s_ = − 0.28, p = 3.6 × 10^−2^, N = 58; Figure S10B). There is no association between CMap 2’s self-retrieval performance and CMap 2-MZS’s within-dataset reproducibility (r_s_ = 0.05, p = 0.69, N = 58; Figure S10C). This indicates there is a relationship between CMap 2’s retrieval performance (when queried with CMap 1 signatures) and within-dataset DE reproducibility.

Since we establish the relationship between CMap 2’s retrieval performance with cross-dataset DE reproducibility (Figure S7), and with within-dataset DE reproducibility (Figure S10), by inference, we would expect both cross-dataset and within-dataset DE reproducibility to also be mutually related. In Figure S11, we see that cross-dataset and within-dataset DE reproducibility is strongly associated for both CMap 1 (r_s_ = 0.59, p = 8.3 × 10^−8^, N = 72; Figure S11A) and CMap 2-MZS (r_s_ = 0.38, p = 2.4 × 10^−3^, N = 61; Figure S11B). With this, we clearly establish a chain of associations, linking CMap 2’s compound retrieval performance with both cross-dataset and within-dataset DE agreement; the latter two are also mutually related. We also establish that the DE “strength” (i.e. number of DE genes in a profile/signature) is associated with both CMap 2’s retrieval performance, and within-dataset DE agreement.

### Compound concentration and cell line background affects within-dataset DE agreement

Next, we considered if compound concentration might affect within-dataset DE reproducibility. We performed correlation analysis on the data, limiting to conditions that have multiple replicates and compounds with multiple concentrations (see Methods). For CMap 1, only one compound was eligible: valproic acid tested in MCF7 cells. We observe a positive correlation between reproducibility and concentration, up to a concentration of 500μM (log_10_ concentration = 2.7; Figure S12). This concentration-dependent effect on transcriptional reproducibility is similarly observed in CMap 2 data (Figure 5). In Figure 5A, vorinostat’s reproducibility increases with increasing concentration, regardless of cell line. In Figure 5B and C, we see the same “dose-dependent curve” for MCF7 cell lines (with greater improvement in reproducibility for trichostatin A), while the curve is mostly unchanging in PC3 cell lines; the latter is due to the lack of data at lower concentrations. In Figure 5D, we see a different trend where the MCF7 cell line treated with wortmannin does not show a strong dose-dependence, whereas the PC3 cell line with a more limited concentration range shows otherwise. This suggests that reproducibility of cellular transcriptional response is dose-dependent and cell line-dependent. In cases where a compound’s transcriptional change appears irreproducible, there are a few possible explanations: either the concentration range tested was too narrow to observe a trend, or the compounds do not induce a strong and consistent cellular transcriptional response at any reasonable concentration.

**Figure 5:**
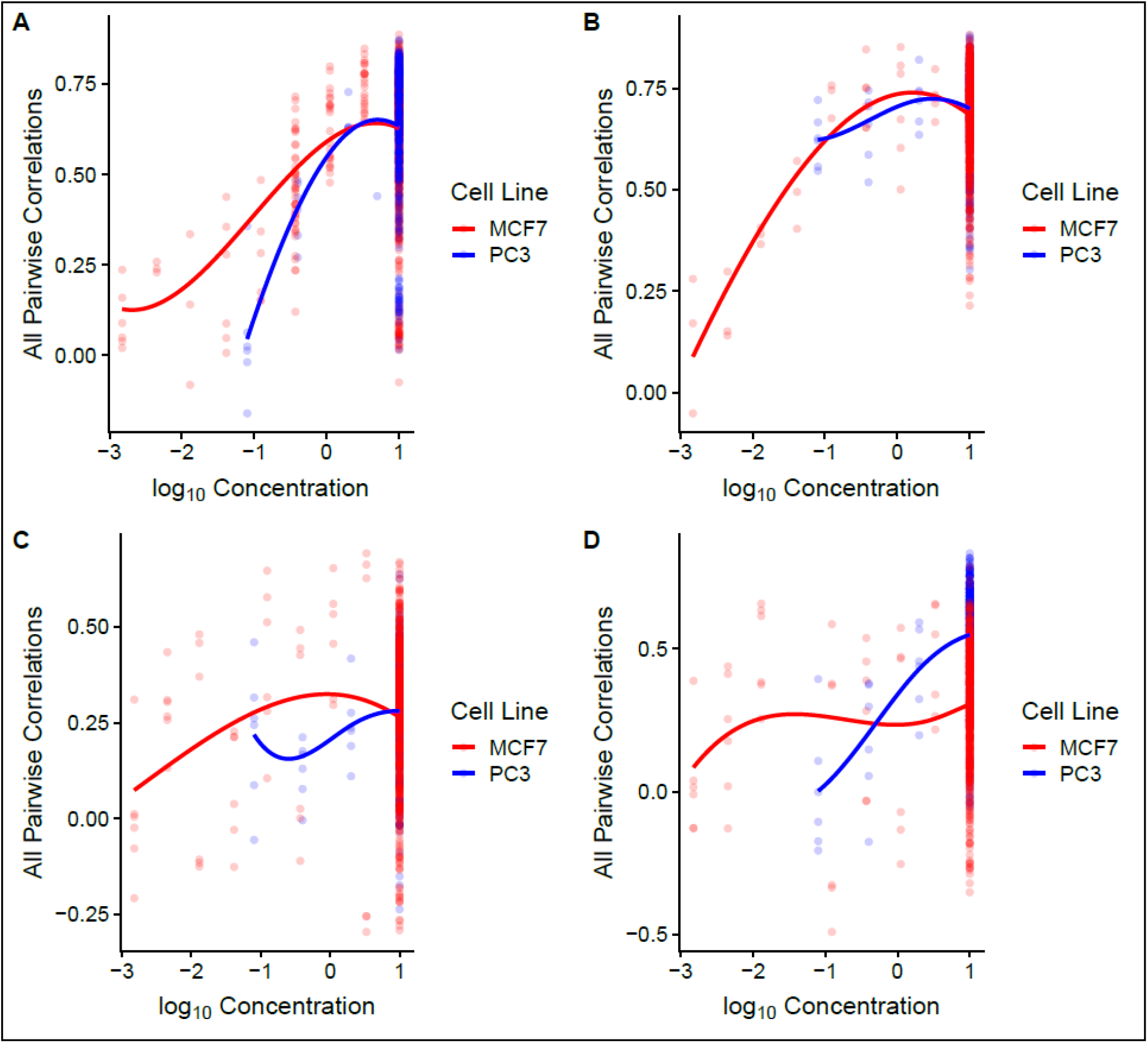
Scatter-plot of all pairwise rank correlations of DE profiles against compound concentration, for four different compounds (vorinostat, trichostatin A, geldanamycin and wortmannin for A-D respectively). Underlying data used is CMap 2-MZS and all genes were used in the calculations. Lines are LOESS fit of the correlations, while colours distinguish the cell lines being tested.

We next considered if both compound concentration and cell line effects are influencing DE reproducibility through DE strength itself. For the same four compounds, in Figure S13, we observe that most compounds have an increasing number of DE genes with increasing compound concentration. It is striking that the trend-lines observed in both Figure 5 and S13 are similar. Put in context with our findings up to this point, we can conclude that in order for a compound to have a reproducible transcriptional change, the transcriptional change needs to be strong, and that is influenced by two factors: appropriate compound concentration and cell line responsiveness.

### CMap 2 has limited within-dataset reproducibility of sample level gene expression

The next angle we considered for CMap data quality is the reproducibility of the underlying expression data at the sample level. We hypothesize that if a pair of DE profiles had such low agreement as seen in Figure 4, the agreement between the constituent samples’ expression profiles used in the two DE replicates would be similarly poor. As we did not work with raw expression data for the CMap 1 dataset (we used pre-computed DE data from Broad Institute), our analysis of sample-level gene expression reproducibility is performed only for CMap 2. Two measures of sample gene expression agreement based on rank correlation (“in-replicate” and “cross-replicate”) were used (see Methods).

In Figure 6, we observe that for both CMap 2-MZS and CMap 2-FC, the distribution of the “in-replicate” agreement of sample expression profiles for the “treatment” group has a mode of 0.95 (with a mean of 0.92), while the “control” group of CMap 2-FC has a mode of 0.96 (with a mean of 0.93). At first glance, this suggests that the sample agreement of gene expression is fairly good. However, this is lower than the typical median correlation of gene expression (r_s_ = 0.98) reported when comparing DE replicates across laboratories (Chen et al., 2007). This observation remains unchanged when we use the maximal “in-replicate” sample agreement (Figure S14) or when the “in-replicate” sample agreement is calculated using landmark genes only (Figure S15). Similar trends are observed when we use the “cross-replicate” sample agreement measure, with the exception that the “control” samples now have distribution properties similar to that of the “treatment” samples (Figure S16). Taken together, it is evident that within-dataset reproducibility of sample gene expression in CMap 2 is lower than commonly observed.

**Figure 6:**
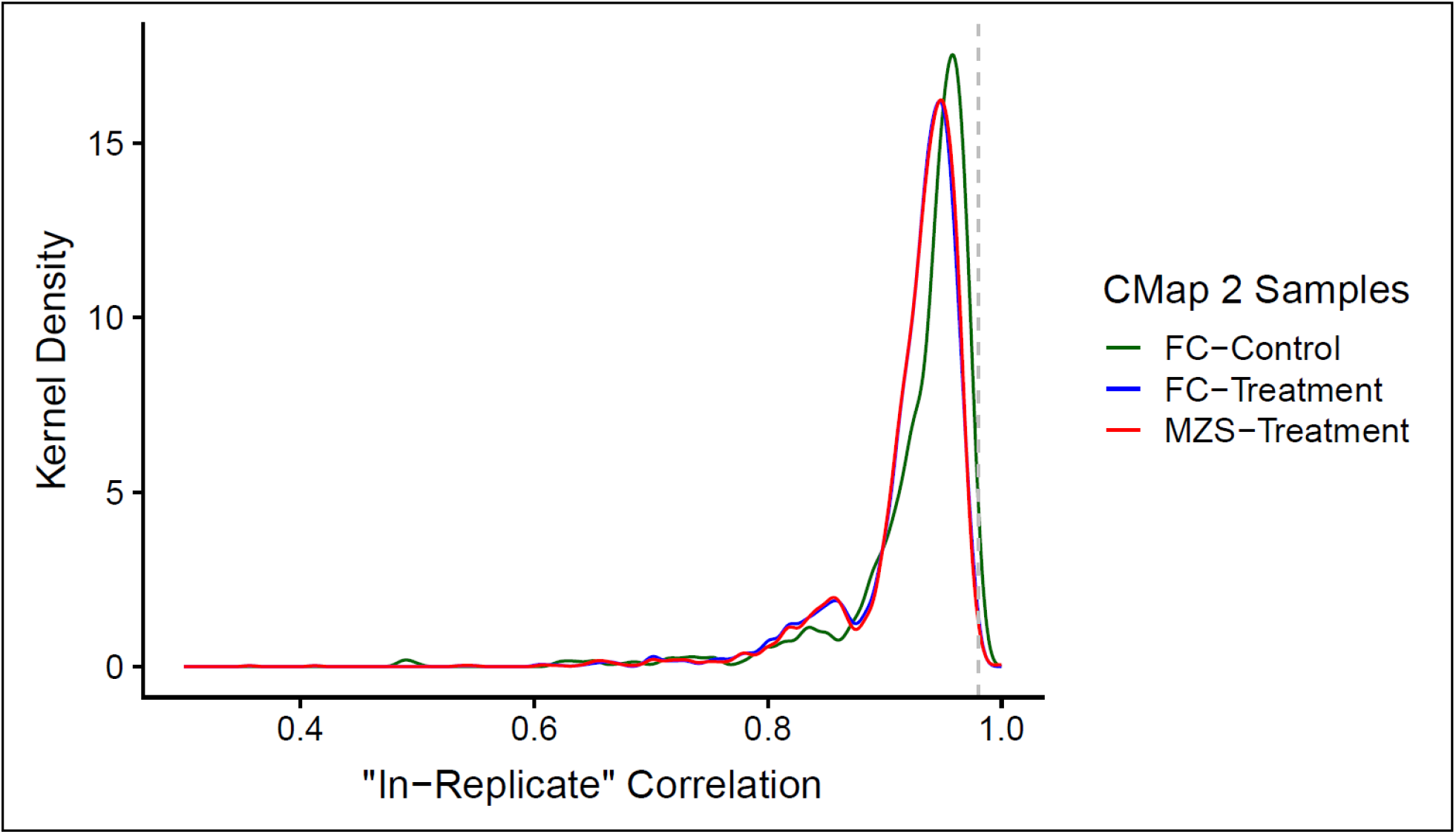
Distribution of rank correlations of “in-replicate” sample gene expression for the DE replicate pairs used in Figure 4, calculated using all genes. The number of unique conditions/pairs assessed for CMap 2-FC and CMap 2-MZS are 2416 and 2477 respectively. The dotted grey line indicates the typical median rank correlation of sample gene expression for intra-platform cross-laboratory comparisons reported in Chen et al. (2007, r_s_ = 0.98).

Next, we assessed if within-dataset agreement of sample gene expression is associated with the number of DE genes, since within-dataset DE agreement is predicted by the number of DE genes (Figure S9; r_s_ = 0.32 for CMap 2). In Figure S17, we see that both “in-replicate” and “cross-replicate” measures of sample-level agreement are lowly correlated with the number of DE genes (r_s_ = −0.29 and −0.22 for “in-replicate” and “cross-replicate” respectively), though not as high as that for within-dataset DE agreement. This indicates that sample-level reproducibility of gene expression is associated with the number of DE genes.

Finally, we considered if sample-level gene expression reproducibility is predictive of within-dataset DE reproducibility. We found that both “in-replicate” and “cross-replicate” measures of sample-level reproducibility to be weakly associated with within-dataset DE reproducibility (r_s_ = 0.16 and 0.18 for “in-replicate” and “cross-replicate” respectively; Figure S18). Considering that the number of DE genes is a fairly strong driver of DE reproducibility, we hypothesized that the weaker signals of association could be teased out using an appropriate statistical model. We build multivariate linear models to test if within-dataset DE agreement is influenced by three different factors: number of DE genes, “in-replicate” agreement of sample expression and “cross-replicate” agreement of sample expression (see Methods for model expression). We found that all three factors are predictive of within-dataset DE agreement (adjusted-R^2^ = 0.372, F (3, 2473) = 490.2, p < 1 × 10^−10^); with the predictive power being statistically significant (Β = 0.60, 0.13, 0.19 for number of DE genes, “in-replicate” and “cross-replicate” agreement of sample expression respectively; p < 1 × 10^−10^ for all three predictors). Taken together, we establish that within-dataset DE agreement is predominantly influenced by the number of DE genes (which in turn is influenced by compound concentration and cell type responsiveness), and less so by sample-level reproducibility of gene expression.

### Neither CMap shows high agreement with third-party profiles

Due to the overall poor agreement between CMap 1 and 2, the final question is raised as to which is more reliable. In an attempt to resolve this question, we compared both CMap datasets to a third dataset, GSE69845 (De Abrew et al., 2016) (referred to as the “DeAbrew dataset”) in which MCF7 cells were exposed to compounds in a similar manner to CMap. Twelve of the compounds in DeAbrew are represented in the harmonized CMap data under comparable conditions (see Methods; Table S4). We found that for 11 of the 12 compounds, the DeAbrew profiles were poorly correlated with both CMaps, similar to the comparison of the two CMaps with each other (mean r_s_ < 0.08); limiting the analysis to only landmark genes gave similar results (mean r_s_ < 0.1). The exception was the profile for vorinostat, regardless whether all genes (r_s_ = 0.38 – 0.72; Figure S19) or only landmark genes (r_s_ = 0.57 – 0.79) are used. As this was the only compound that had high agreement between CMap 1 and 2, it does not resolve the question of which CMap dataset is more reliable.

## Discussion

In this paper, we show that drug prioritizations obtained with CMap 1 and 2 are rarely concordant. Our investigation shows that both datasets, at the DE level, suffer from data reproducibility issues both *within* and *between* the two CMaps. We also demonstrated, through a chain of associations, that a profile’s DE “strength” (number of DE genes) and sample-level reproducibility of gene expression are predictors of DE reproducibility and compound retrieval performance of CMap. We found that DE strength is influenced by both compound concentration and cell-type responsiveness. Furthermore, our attempt to identify which of the two CMaps is “better” was unsuccessful, as the third dataset we compared to also show poor agreement with both CMaps. Our results lead to a number of recommendations for users of the current CMaps, and can guide future development and evaluation of drug repositioning resources.

The clearest previous indication that CMap results might not be reproducible came from the study of De Abrew et al. (2016), in which they compared their CMap-like data to CMap 1 (their work predated CMap 2). We stress that their interpretation was much more positive. However, their analysis was qualitative, and they used relaxed criteria to judge agreement. Thus while it is true that there is an overall trend of positive “Connectivity Scores” when querying CMap 1 using De Abrew’s dataset, the actual rankings are not in agreement, and only vorinostat and vinblastine had concordant retrieval performance. In actual use, investigators are unlikely to look very far down the list of recommended compounds, so agreement should be judged by the compound being ranked very highly, e.g. within the top-20 at worst. By this criterion, De Abrew’s findings are in clear agreement with ours, and we further show that the De Abrew data does not agree well with CMap 2. De Abrew et al. also noted that the qualitative agreement of their results with CMap 1 was higher for the highest concentration of compounds. Our results confirm this finding with a major caveat: DE reproducibility is indeed concentration-dependent (Figure 5); however the highest compound concentrations do not necessarily guarantee high cross-dataset reproducibility/retrieval (Figure 1). In fact, a major predictor of profile reproducibility is the DE strength of a profile, and this factor is influenced by both compound concentration and cell type responsiveness (Figures S9 and S13); the other predictor being the reproducibility of the profiles’ constituent samples’ gene expression (Figure S18). We observe that if the concentration is in the incorrect range to cause strong DE or if a cell type does not respond to the tested compound, it is unlikely to yield a reproducible signature, and thus lead to poor prioritization results. The low DE strength is also likely to explain why we observe poor profile reproducibility across the three datasets (CMap 1, CMap 2 and De Abrew).

Related to these observations, we raise a concern about limitations of the Connectivity Map methodology. The CMap concept is based on the similarity/enrichment of transcriptional change between profiles. We observe that this method is likely to be successful only if the transcriptional change of the profiles (be it in the query or the corpus) are both strong and reproducible. From our observations in Figure 4, it could be postulated that a fair portion of the current CMap corpus may not be very useful in its current state. Interestingly, in order to reduce the distorting influence of outlier samples, the LINCS team employ a weighting mechanism during the process of generating their DE data (i.e. Level 5 data). Our observations in Figures 6 and S18 suggest that the weighting mechanism may not be effective, especially facing data where the sample-level gene expression data itself are irreproducible to begin with.

Our findings raise the question then of how investigators have been able to successfully apply CMap, inasmuch there are positive findings in which some follow-up experiment yielded encouraging results in both CMap 1 and 2 (see Introduction; CMap 2 was published more recently so there are fewer such reports). If we interpret these reports to indicate that both CMap 1 and 2 “work”, it seems mysterious that the two datasets usually give different results for the same compounds, and the De Abrew data only reinforces this conclusion. One potential explanation is that we were very unlucky and the ~600 compounds we were able to compare across the datasets are highly enriched for poor performers. Another possibility is that both CMaps do recommend valid compounds – just different ones. However, one would still expect that CMap 2 would highly rank the queried compound (relative to other CMap 1 compounds) when probed with its CMap 1 signature, and that was usually not the case (Figures 1 and S1). In our view, the literature is not necessarily inconsistent with our findings. First, there may be a “file drawer effect”, in that negative results (i.e. failure of validation) obtained with either CMap would tend to be published at a lower rate than successes. Second, despite over ten years passing since CMap 1 was released, we have been unable to identify cases where it has been used to actually reposition a drug for extensive research or clinical use. Our survey of the literature suggests that for most CMap success stories, there are no follow-up confirmatory studies after the initial report.

Based on these observations and considerable uncertainty about which, if any, version of CMap is reliable, we make some recommendations for researchers. Obviously, caution is advised and stringent follow-up validation experiments should be performed on any candidate compounds. More importantly however, investigators should give thought to the relevance of the cell lines used in CMap to their studies. Drug effects are likely to be cell-type specific (Figure 5), and indeed this is the motivation for CMap using multiple cell lines. However, most users query CMap with signatures obtained from cell types or tissues not used in CMap. The fact that querying CMap 2 with CMap 1 signatures of the same conditions gives poor results (Figures 1 and S1) raises the question of how querying CMap with a signature from a heterologous cell type would yield relevant results. In fact, the LINCS team have reported only 26% of compounds having a similar signal across cell lines (Subramanian et al., 2017), indicating that querying LINCS with a non-Touchstone cell line signature may not produce usable results. If possible, generating a signature using one of the cell lines included in CMap would be a sound approach to mitigate this concern, e.g. (Johnstone et al., 2012). Next, for users of CMap 2 (LINCS) in particular, we have two recommendations: First, if reasonable, they could use query lists with more genes, since it appears to provide slightly better compound retrieval performance (Figures S2 and S3); and second, they can filter out (or assign lower confidence to) some of the recommended compounds if CMap 2 has poor self-retrieval performance of those compounds (Figure S4). Separately, if possible both CMap 1 and 2 should be queried and the results compared; concordant results might be viewed favorably, or perhaps all compounds recommended by either CMap should be considered for follow-up. Finally, if pursuing wet-lab validation, it would be wise to conduct control experiments with compounds that were not ranked highly, in addition to ones which are highly ranked, e.g. (Ryals et al., 2018). If the CMap recommendations are valid, the lowly-ranked compounds should validate at a much lower rate than the highly ranked ones.

## Acknowledgments

We thank Ogan Mancarci and German Novakovsky, who initially brought the question of reproducibility of CMap to our attention. We also thank Aman Sharma, Calvin Chang and Owen Tsai for assistance with data curation. This work was supported by grants from the National Institute of Health (MH111099), the Natural Sciences and Engineering Research Council of Canada (RGPIN-2016-05991) and a University of British Columbia Four-Year Doctoral Fellowship to N.L.

## Author Contributions

P.P. and N.L. designed the study; N.L. performed data collection and analysis; P.P. and N.L. wrote the manuscript.

