## Supplementary material for "Evaluation of Connectivity Map shows limited reproducibility in drug repositioning": Supplements.pdf

**Table S1:** Combinations of compounds, concentrations and cell lines that are common between both CMap 1 and 2 (“Harmonized data”). The treatment duration for all combinations is fixed at 6 hours.

**Table S2:** Combinations of compounds and cell lines used as signatures from CMap 1 to perform L1000-Query on CMap 2 data; strongest compound concentrations are used. The treatment duration for all combinations is fixed at 6 hours and all compounds are part of CMap 2’s Touchstone data subset.

**Table S3:** Combinations of compounds and cell lines used as signatures from CMap 1 to perform L1000-Query on CMap 2 data. The treatment duration for all combinations is fixed at 6 hours and all compounds are part of CMap 2’s Touchstone data subset. Unlike Table S2, the signatures used in Table S3 are from the harmonized data. For the “Hybrid Threshold” case (see Methods), the signatures “10μM rosiglitazone in MCF7” and “10μM scopolamine in PC3” were excluded due to inadequate number of differentially expressed genes.

| Chemical Compound | CMap 1 Concentration (µM) | CMap 2 Concentration (µM) | De Abrew et al. Concentration (µM) |
|---|---|---|---|
| GENISTEIN | 10.0 | 10.0 | 10.0 |
| METFORMIN | 10.0 | 10.0 | 10.0 |
| PHENFORMIN | 10.0 | 10.0 | 10.0 |
| TROGLITAZONE | 10.0 | 10.0 | 10.0 |
| VORINOSTAT | 10.0 | 10.0 | 10.0 |
| CHENODEOXYCHOLIC-ACID | 10.2 | 10.0 | 10.0 |
| GRISEOFULVIN | 11.2 | 10.0 | 10.0 |
| METHOTREXATE | 8.8 | 10.0 | 10.0 |
| CLOBETASOL | 8.6 | 10.0 | 10.0 |
| KETOCONAZOLE | 7.6 | 10.0 | 10.0 |
| PROGESTERONE | 12.8 | 10.0 | 10.0 |
| FLUTAMIDE | 14.4 | 10.0 | 10.0 |

**Table S4:** Compounds and concentrations used in the comparison of CMap 1 and 2 against the De Abrew dataset. The treatment duration for all compounds is fixed at 6 hours and the cell line used is MCF7. With the exception of the first five compounds (genistein, metformin, phenformin, troglitazone and vorinostat), we chose conditions that have a compound concentration of between 5 - 15µM in CMap 1 (as the corpus lacks similar conditions with exactly 10µM).

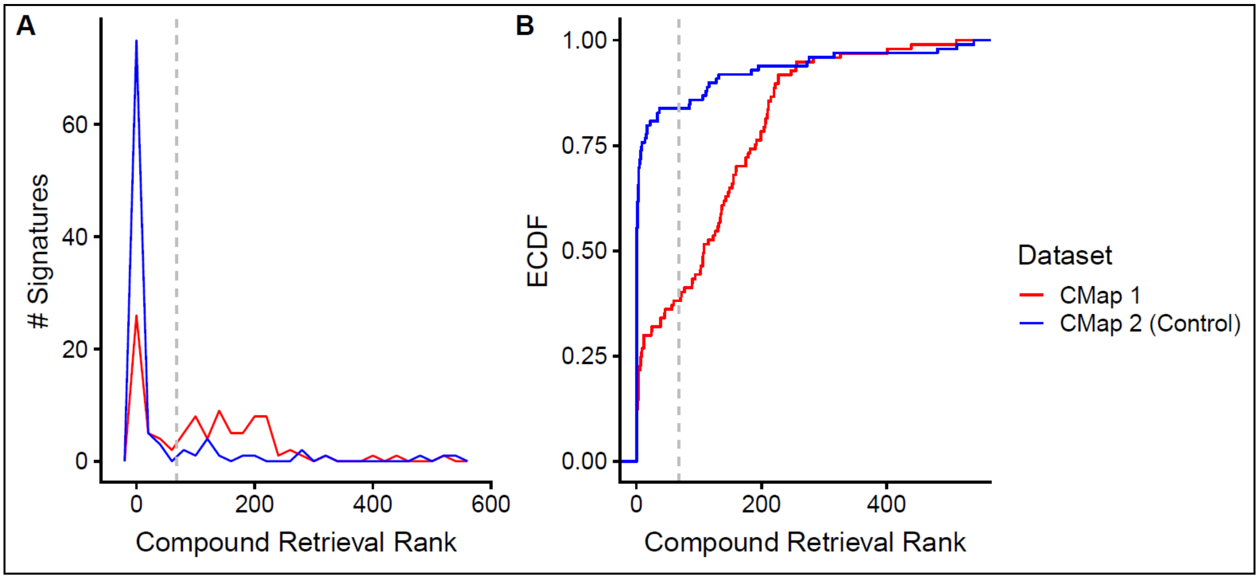


**Figure S1:** Distribution of compound retrieval ranks (1 is best) from querying CMap 2 using signatures derived from CMap 1 (red lines; “Hybrid Threshold”) or CMap 2 (blue lines; self-query) data, as in main Figure 1, but with the added constraint that the conditions of the signatures are from the “harmonized data” subset (Table S3, N = 97). A shows the distribution while B is the empirical cumulative distribution function (ECDF) of the same results. The dotted grey line indicates the profiles where the query compound is ranked in the top 10% (rank ⩽ 68).

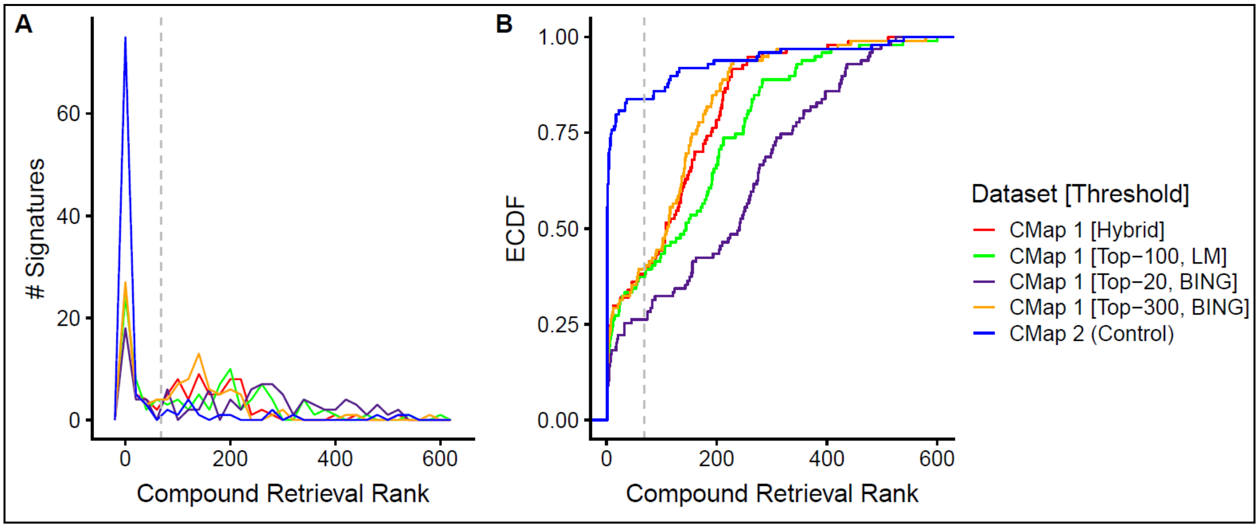


**Figure S2:** Distribution of compound retrieval ranks (1 is best) from querying CMap 2 using signatures derived from CMap 1 (non-blue lines) or CMap 2 (blue lines) data, as in Figure S1; the various CMap 1 signatures are derived using different thresholding methods (See Methods; Table S3, N = 97). A shows the distribution while B is the empirical cumulative distribution function (ECDF) of the same results. The dotted grey line indicates the profiles where the query compound is ranked in the top 10% (rank ⩽ 68).

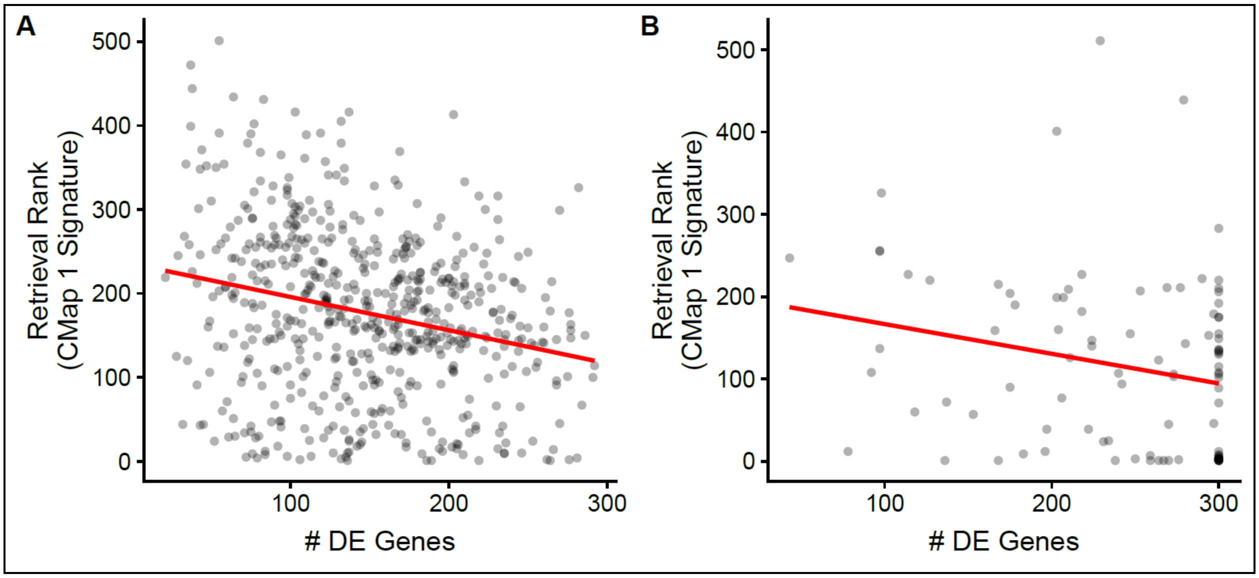


**Figure S3:** Scatter-plot of compound retrieval ranks from querying CMap 2 using signatures derived from CMap 1 data against total number of DE genes (in the CMap 1 signatures). Linear regression of the data is shown in red. The underlying data in A is from the 588 query signatures of the strongest concentration (Table S2); B is from the 97 query signatures from the harmonized data (i.e. common concentration) using the "Hybrid Threshold" (Table S3). The Spearman correlations between retrieval ranks and total number of DE genes are -0.24 and -0.26 for the data in A and B respectively.

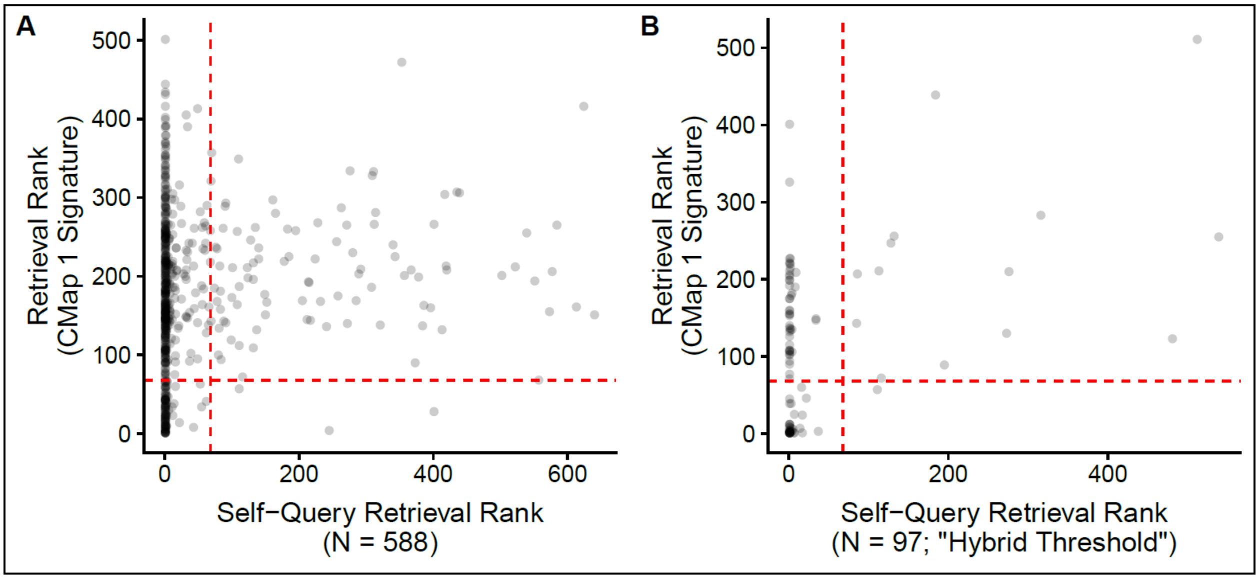


**Figure S4:** Scatter-plot of compound retrieval ranks from querying CMap 2 using CMap 1 data-derived signatures against those of CMap 2 self-queries. The dotted red lines indicate the profiles where the query compound is ranked in the top 10% (rank ⩽ 68). The underlying data in A is from the 588 query signatures of the strongest concentration (Table S2); B is from the 97 query signatures from the harmonized data (i.e. common concentration) using the "Hybrid Threshold" (Table S3).

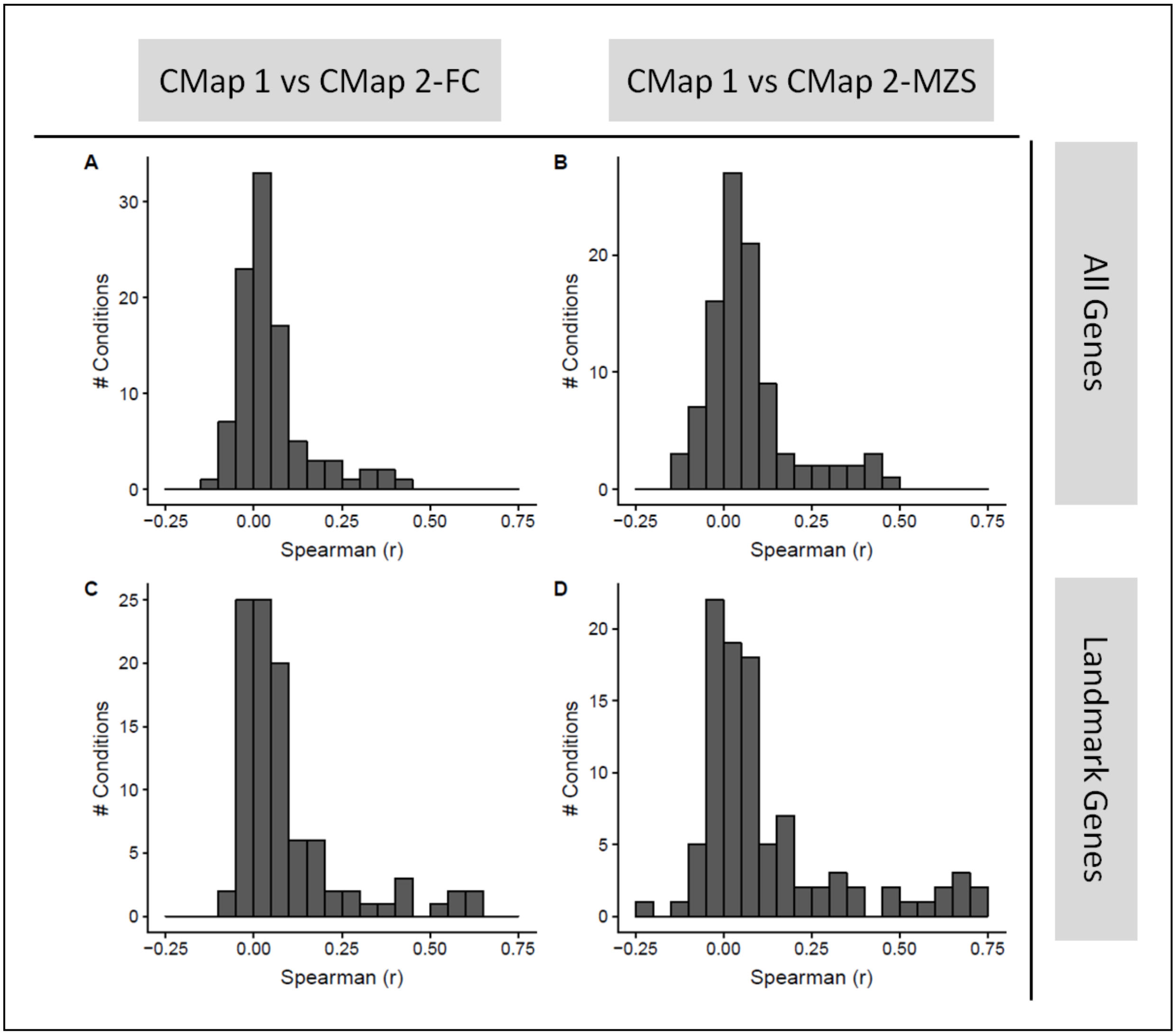


**Figure S5:** Distribution of pairwise rank correlation of DE profiles from CMap 1 against CMap 2; profiles are limited to compounds in the "Touchstone" subset. A and C are comparisons against CMap 2-FC; B and D against CMap 2-MZS. All genes common between CMap 1 and CMap 2 are used in A and B, while only "landmark genes" are used in C and D. (N = 98 conditions).

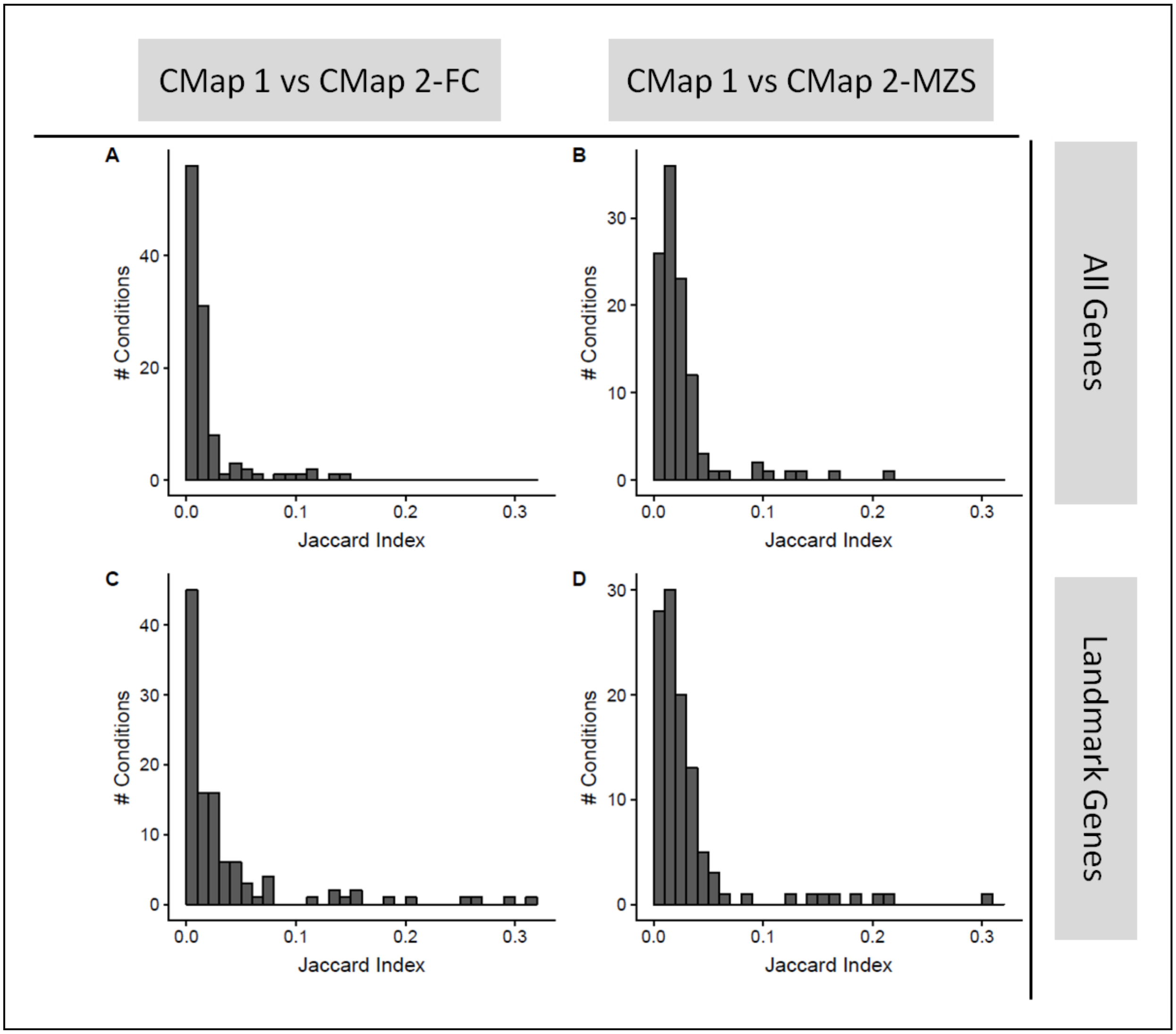


**Figure S6:** Distribution of pairwise Jaccard index of DE profiles from CMap 1 against CMap 2; underlying data is similar to main Figure 2, but using a different similarity metric. A and C are comparisons against CMap 2-FC; B and D against CMap 2-MZS. All genes common between CMap 1 and CMap 2 are used in A and B, while only "landmark genes" are used in C and D. (N = 109 conditions).

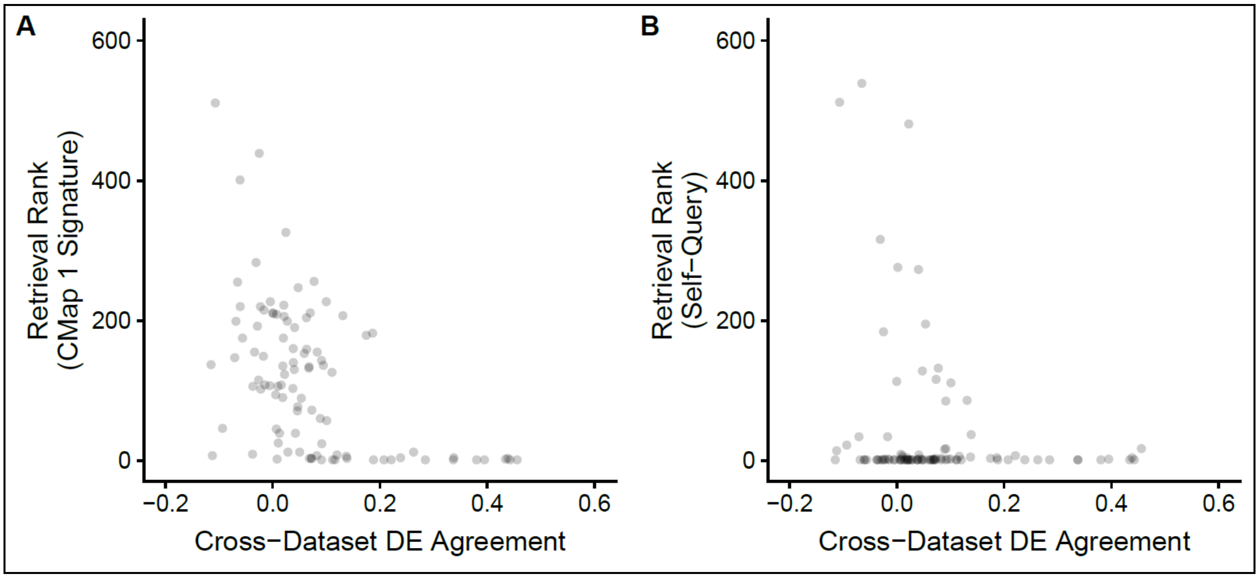


**Figure S7:** Scatter-plot of compound retrieval ranks from querying CMap 2 with signatures derived from CMap 1 or CMap 2 data (y-axis; A and B respectively) against cross-dataset DE agreement (between CMap 1 and CMap 2-MZS; N = 97); the cross-dataset agreement values reported here are calculated using rank correlation on all genes.

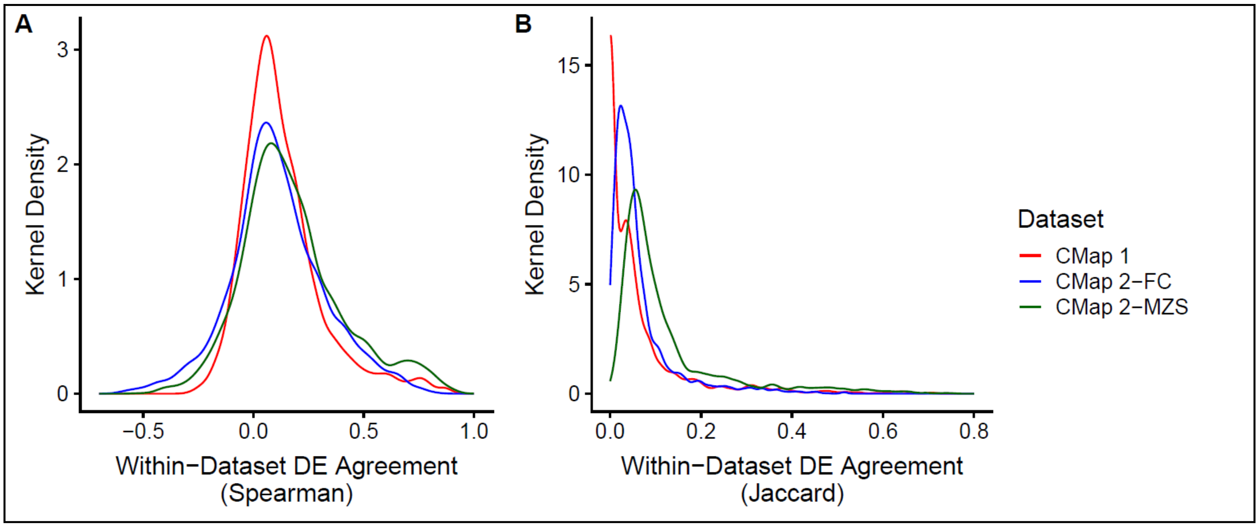


**Figure S8:** Distribution of maximum ("best case") pairwise similarity values (A: rank correlation, B: Jaccard index) for DE profile replicates of the same condition calculated using landmark genes, within each dataset. The number of unique conditions assessed for CMap 1, CMap 2-FC and CMap 2-MZS are 1614, 2416 and 2477 respectively.

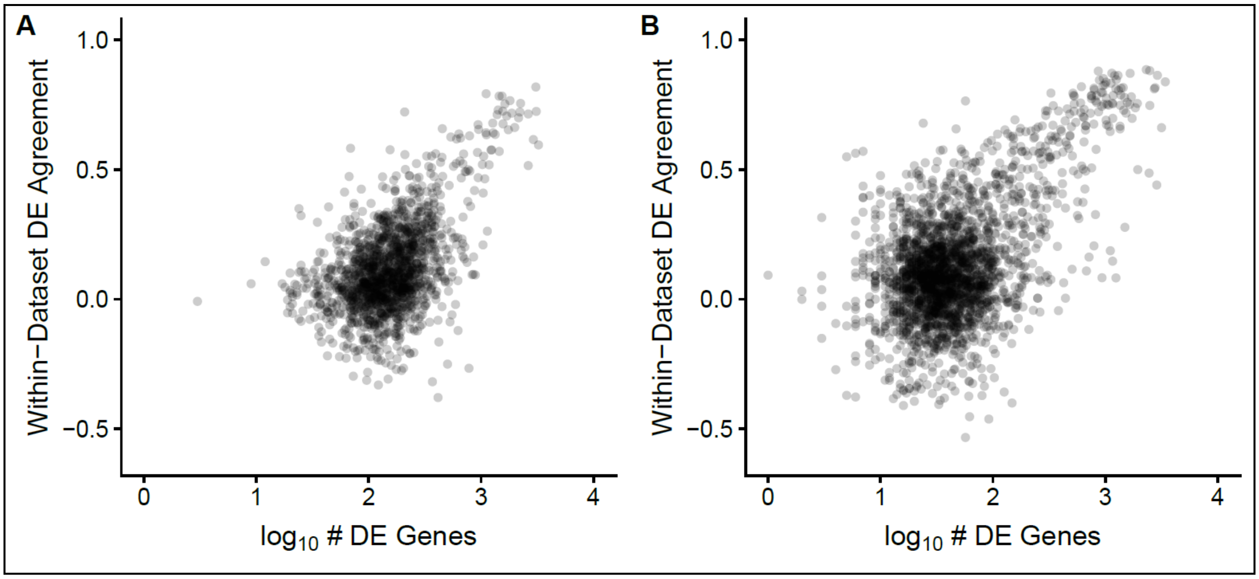


**Figure S9:** Scatter-plot of within-dataset DE agreement (A: CMap 1, B: CMap 2-MZS) against the $\log_{10}$-transformed number of DE genes of the replicate pair; only the maximum ("best case") rank correlations for each unique condition/pair is used; for the number of DE genes, the lesser ("limiting case") of the two replicates in the comparison pair is used. The number of conditions assessed for CMap 1 and CMap 2-MZS are 1614 and 2477 respectively.

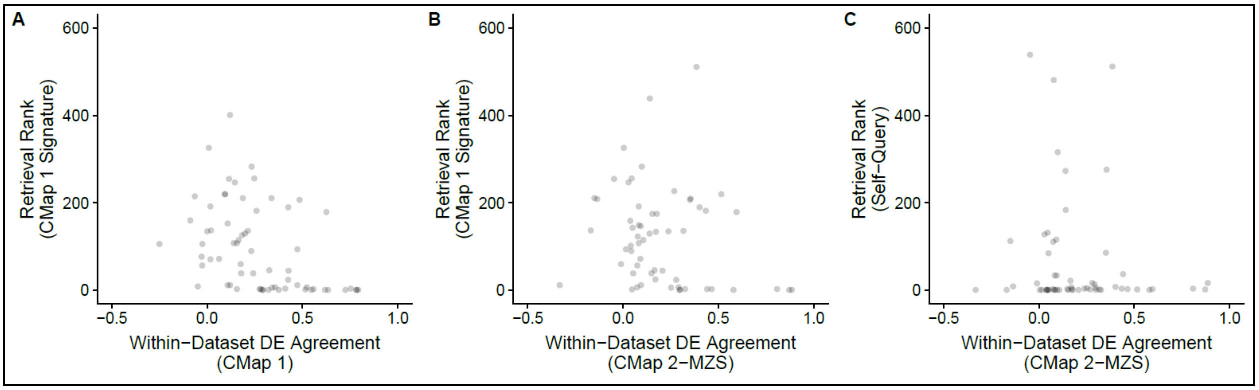


**Figure S10:** Scatter-plot of compound retrieval ranks from querying CMap 2 with signatures derived from CMap 1 (y-axis; A and B) or CMap 2 (C) data against the within-dataset DE agreement (x-axis; CMap 1 for A, CMap 2-MZS for B and C). As with Figure S9, only the maximum rank correlation for each unique condition is shown for the within-dataset agreement. Total number of conditions shown in A is 65; for B and C are 58.

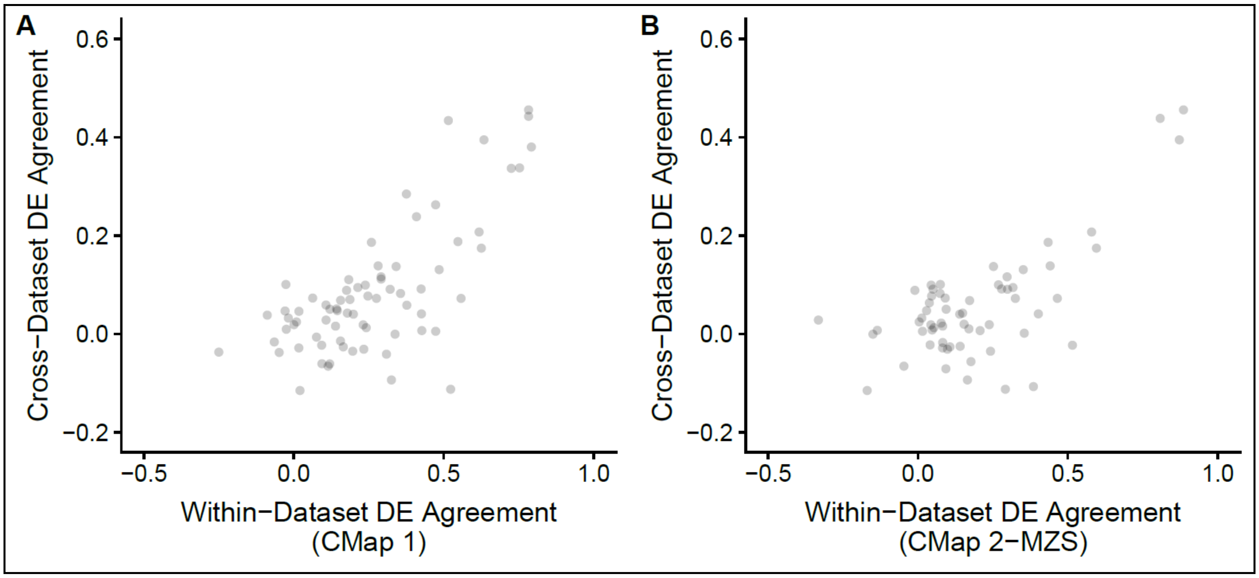


**Figure S11:** Scatter-plot of cross-dataset DE agreement (between CMap 1 and CMap 2-MZS) against the within-dataset DE agreement (A: CMap 1, B: CMap 2-MZS). As with Figure S9, only the maximum rank correlation for each unique condition is shown for the within-dataset agreement. Total number of conditions shown in A and B are 72 and 61 respectively.

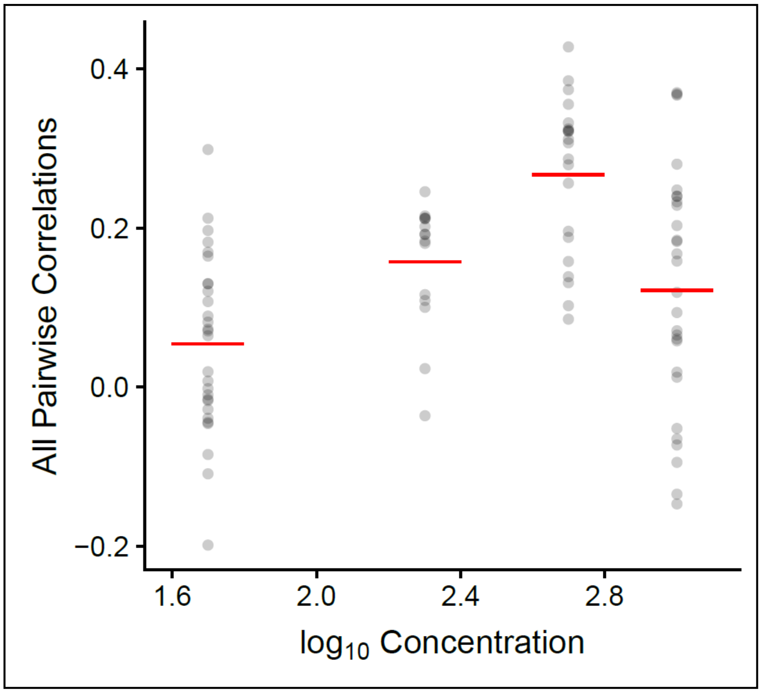


**Figure S12:** Scatter-plot of all pairwise rank correlations of DE profiles against compound concentration, for valproic acid. Underlying data used is CMap 1, treated cell line is MCF7 and all genes were used in the calculations. Red lines indicate the mean correlation for each concentration group.

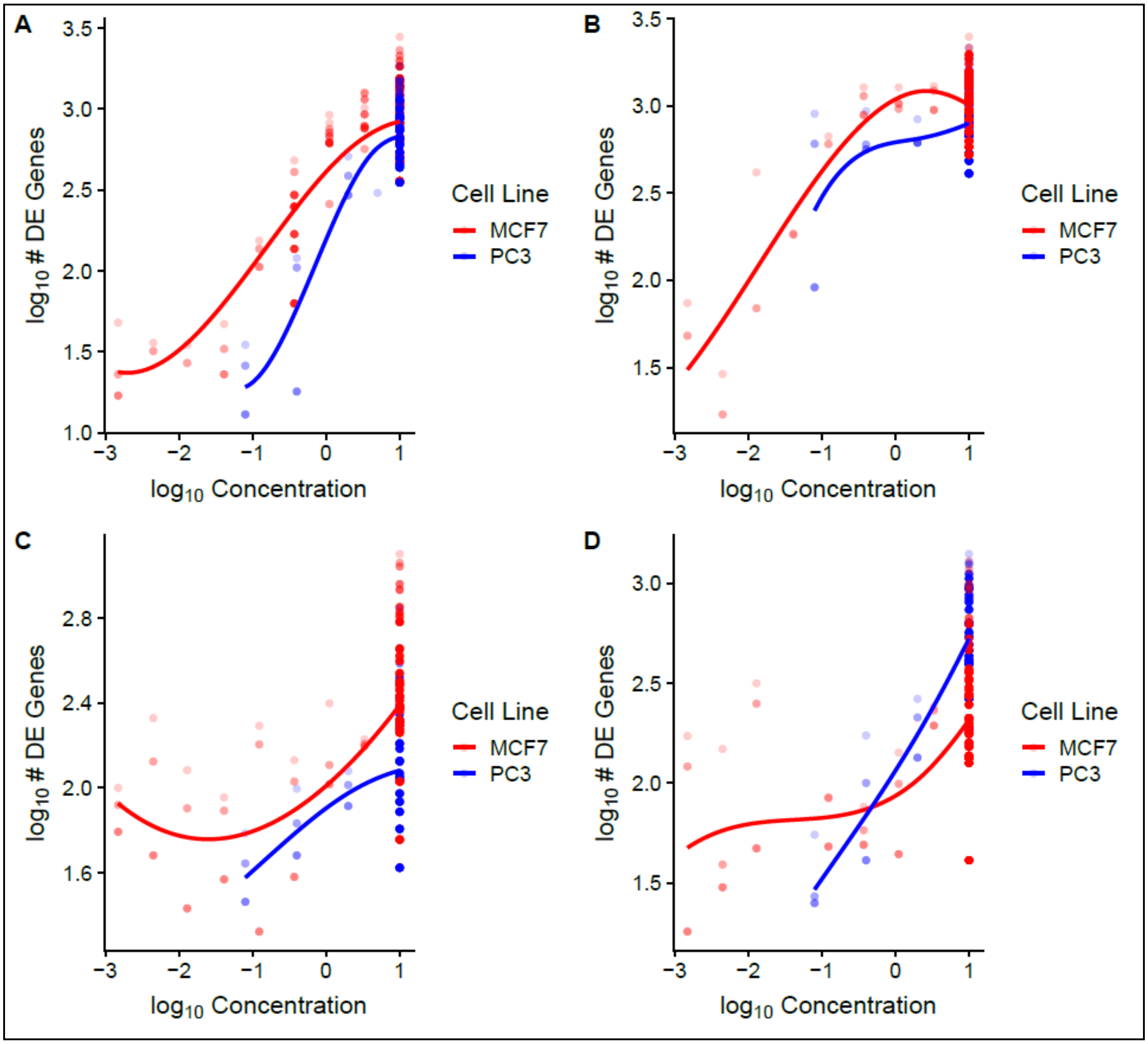


**Figure S13:** Scatter-plot of DE “strength” of profile replicate pairs against compound concentration, for four different compounds (vorinostat, trichostatin A, geldanamycin and wortmannin for A-D respectively); DE “strength” refers to the $\log_{10}$-transformed number of DE genes of the replicate pair (as used in Figure S9). Underlying data used is CMap 2-MZS and all genes were used in the calculations. Lines are LOESS fit of the values, while colours distinguish the cell lines being tested.

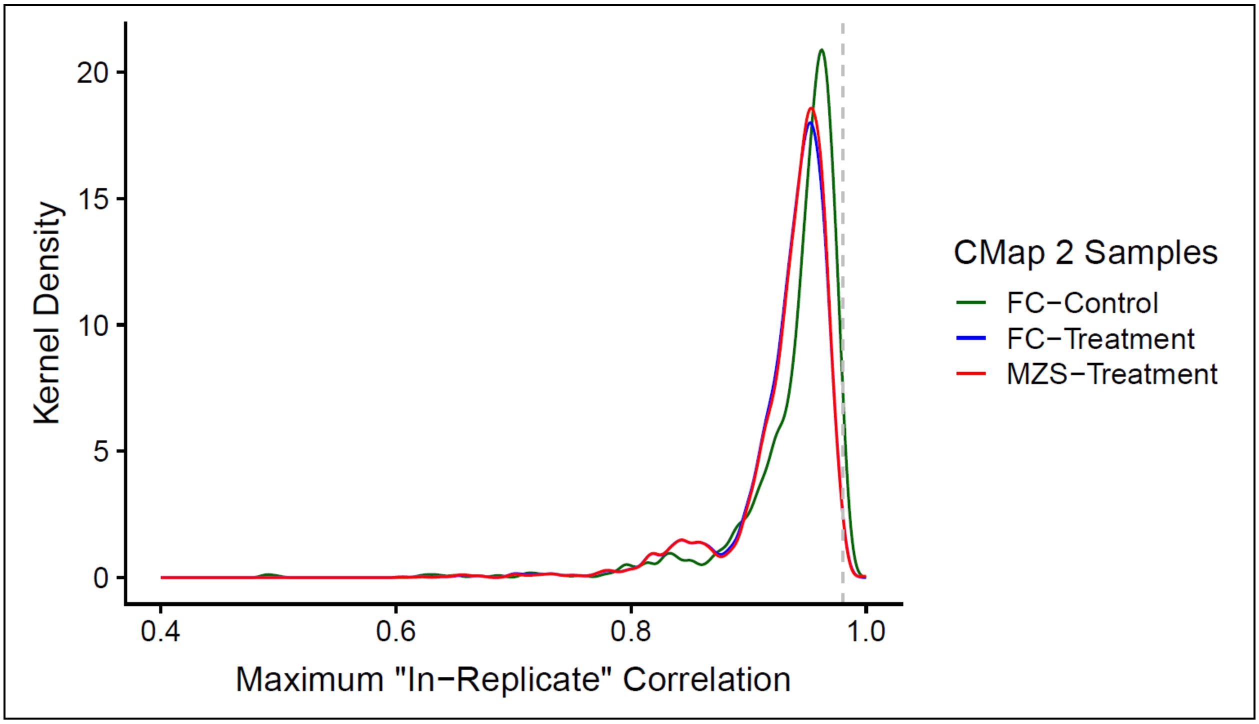


**Figure S14:** Distribution of rank correlations of maximum “in-replicate” sample gene expression (“best case”) for the DE replicate pairs used in Figure 4, calculated using all genes. Unlike main Figure 6 in which the plotted “in-replicate” sample correlations are from the DE replicate pairs with the highest within-dataset DE agreement for each unique condition (i.e. main Figure 4), the sample correlations plotted here are not constrained by the performance of the DE pair. The number of unique conditions/pairs assessed for CMap 2-FC and CMap 2-MZS are 2416 and 2477 respectively. The dotted grey line indicates the typical median rank correlation of sample gene expression for intra-platform cross-laboratory comparisons reported in Chen et al. (2007, $r_s = 0.98$).

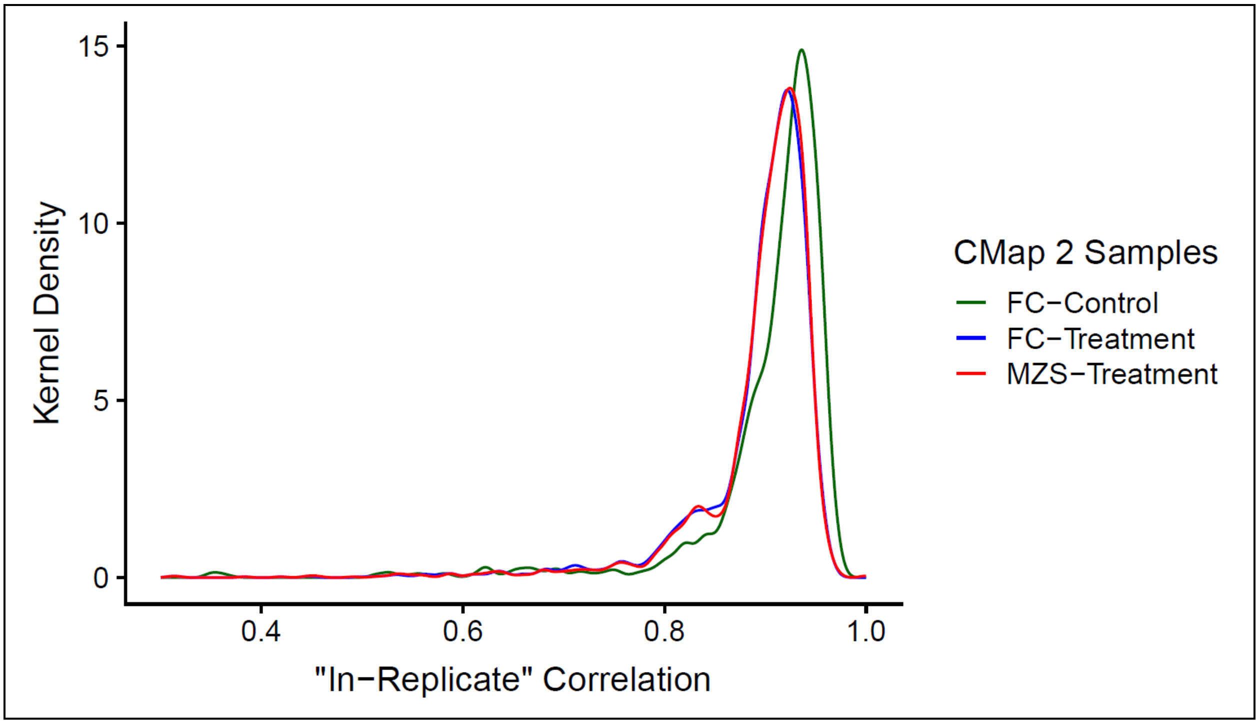


**Figure S15:** Distribution of rank correlations of "in-replicate" sample gene expression for the DE replicate pairs used in Figure 4, calculated using landmark genes only. The number of unique conditions/pairs assessed for CMap 2-FC and CMap 2-MZS are 2416 and 2477 respectively.

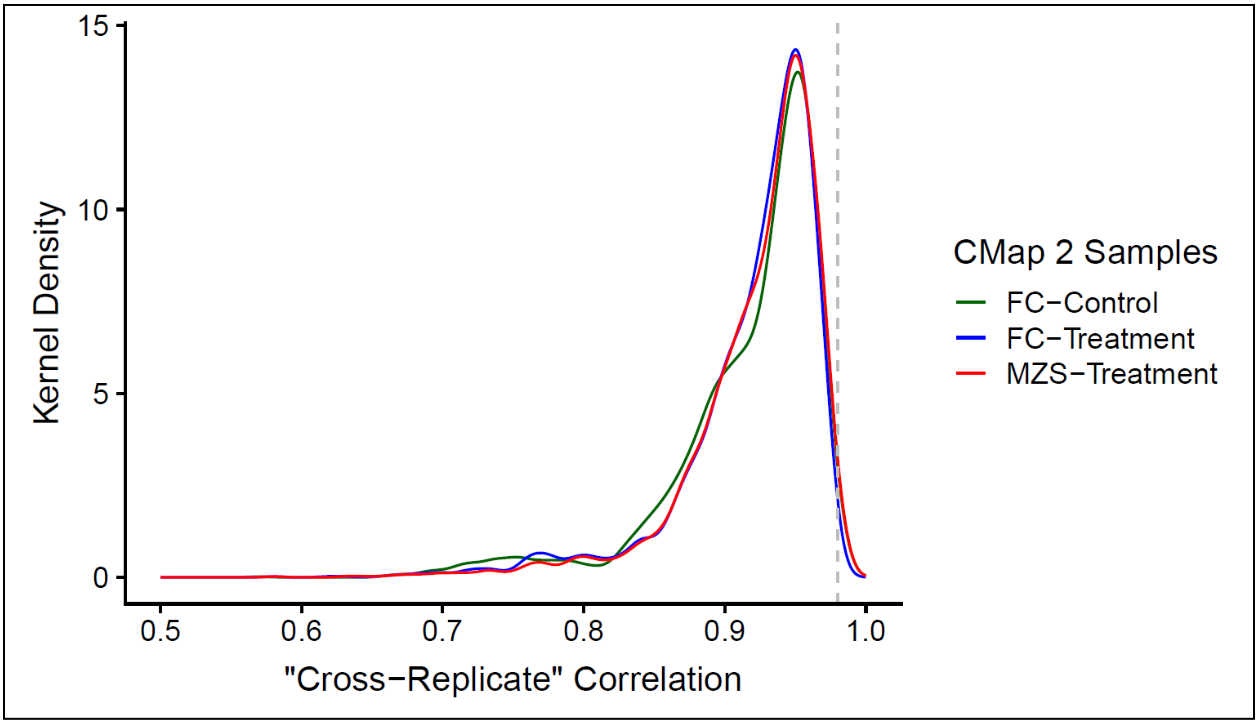


**Figure S16:** Distribution of rank correlations of "cross-replicate" sample gene expression for the DE replicate pairs used in Figure 4, calculated using all genes. The number of unique conditions/pairs assessed for CMap 2-FC and CMap 2-MZS are 2416 and 2477 respectively. The dotted grey line indicates the typical median rank correlation of sample gene expression for intra-platform cross-laboratory comparisons reported in Chen et al. (2007, $r_s = 0.98$).

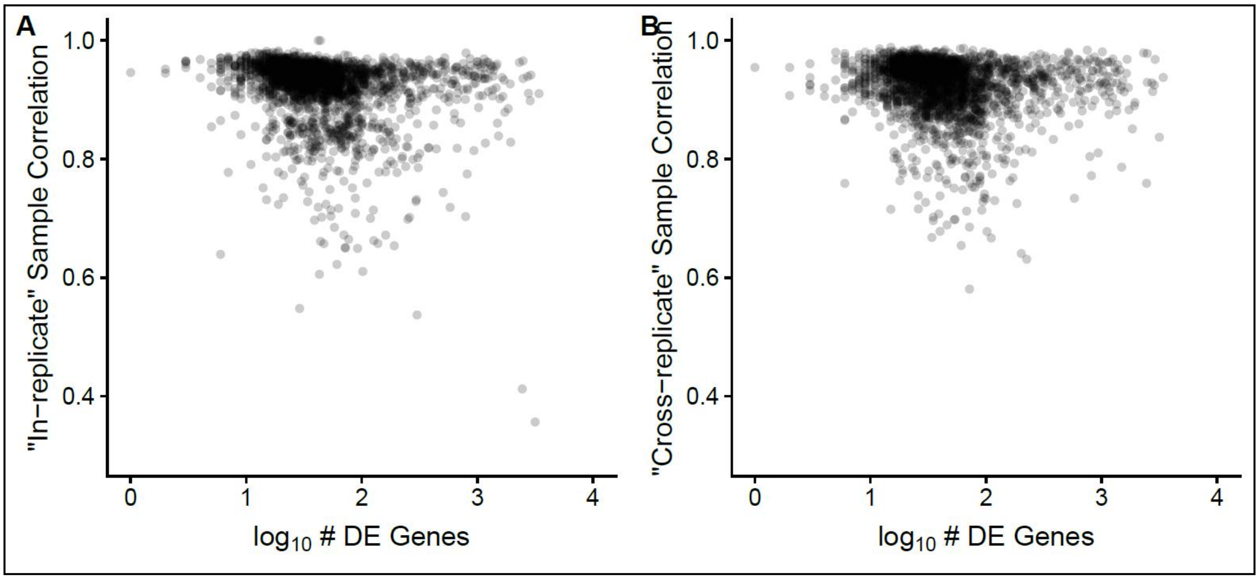


**Figure S17:** Scatter-plot of within-dataset agreement of sample-level gene expression ("in-replicate" and "cross-replicate" for A and B respectively) against the $\log_{10}$-transformed number of DE genes of the replicate pair; as with Figure S9, the lesser number of DE genes between the replicates of a unique condition is used. Underlying data used is CMap 2-MZS and the number of unique conditions assessed is 2477.

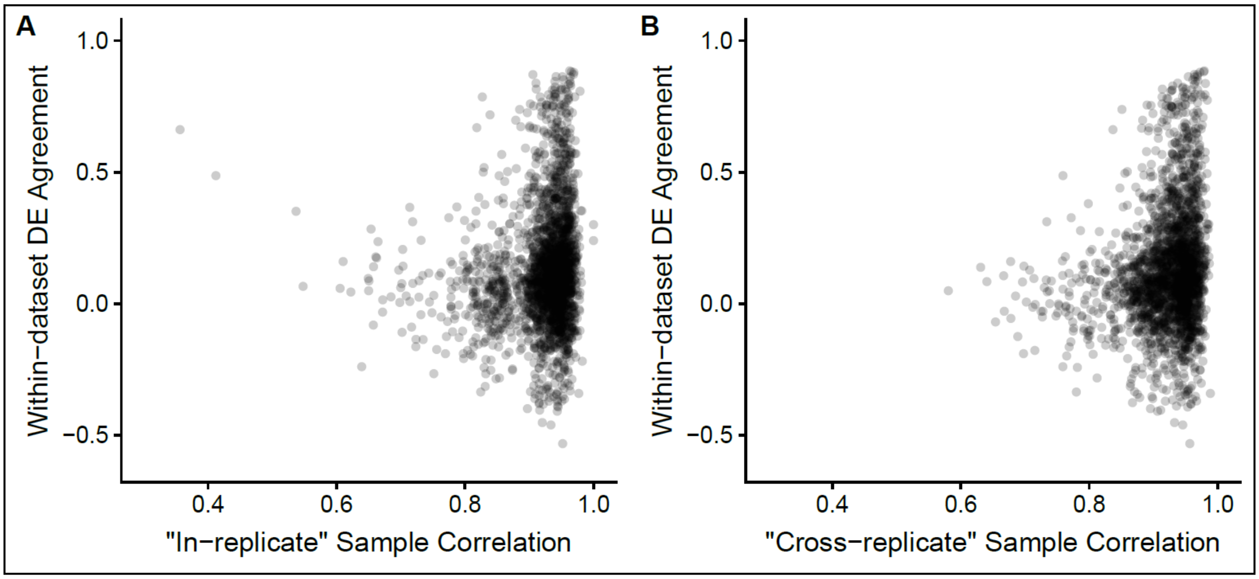


**Figure S18:** Scatter-plot of within-dataset DE agreement against within-dataset agreement of sample-level gene expression ("in-replicate" and "cross-replicate" for A and B respectively). Underlying data used is CMap 2-MZS and the number of unique conditions assessed is 2477.

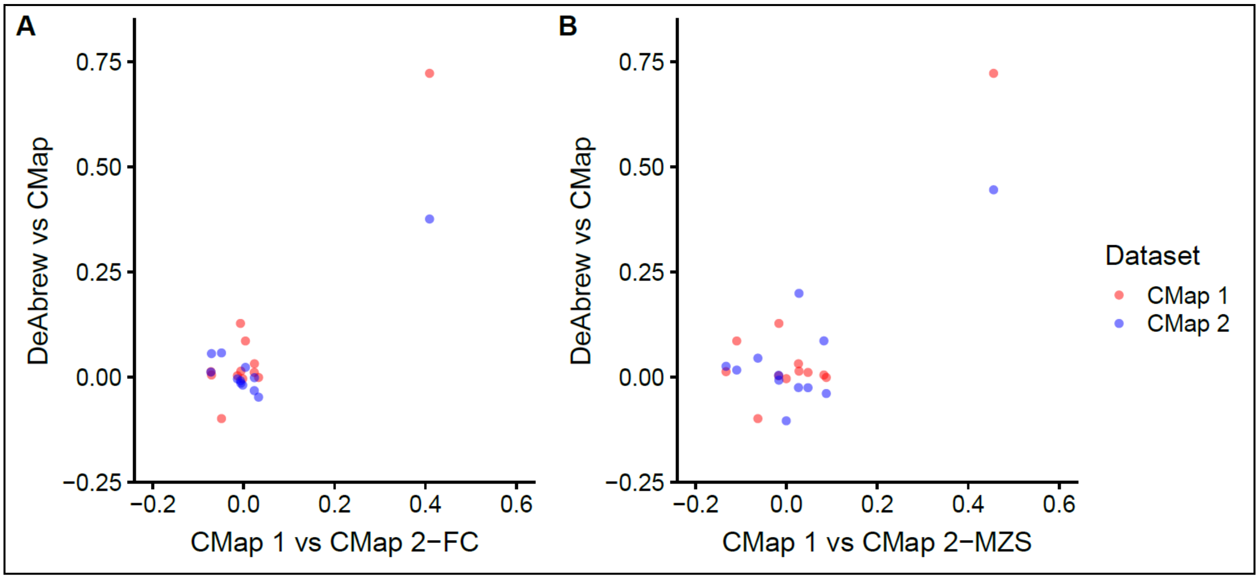


**Figure S19:** Scatter-plot of DE reproducibility between the De Abrew dataset and CMap (y-axis; CMap 1 = red dots, CMap 2 = blue dots) against cross-dataset DE reproducibility between CMap 1 and CMap 2 (x-axis). The underlying versions of CMap 2 used in A and B are CMap 2-FC and CMap 2-MZS respectively. Each data point is 1 of the 12 compounds used to compare between De Abrew to CMap 1 to CMap 2 (Table S4).
